# Structural snapshots uncover a lock-and-key type conserved activation mechanism of β-arrestins by GPCRs

**DOI:** 10.1101/2022.10.10.511556

**Authors:** Jagannath Maharana, Parishmita Sarma, Manish K. Yadav, Sayantan Saha, Vinay Singh, Shirsha Saha, Mohamed Chami, Ramanuj Banerjee, Arun K. Shukla

## Abstract

Agonist-induced phosphorylation of G protein-coupled receptors (GPCRs) is a key determinant for the binding and activation of multifunctional regulatory proteins known as β-arrestins (βarrs). Although the primary sequence and phosphorylation pattern of GPCRs are poorly conserved, the downstream functional responses mediated by βarrs such as receptor desensitization, endocytosis and signaling are broadly applicable across GPCRs. A conserved principle of βarr activation, if any, upon their interaction with different GPCRs harboring divergent phosphorylation patterns remains to be visualized, and it represents a major knowledge gap in our current understanding of GPCR signaling and regulatory paradigms. Here, we present four structural snapshots of activated βarrs, in complex with distinct phosphorylation patterns derived from the carboxyl-terminus of three different GPCRs, determined using cryogenic-electron microscopy (cryo-EM). These structures of activated βarrs elucidate a “lock-and-key” type conserved mechanism of βarr activation wherein a P-X-P-P phosphorylation pattern in GPCRs interacts with a spatially organized K-K-R-R-K-K sequence in the N-domain of βarrs. Interestingly, the P-X-P-P pattern simultaneously engages multiple structural elements in βarrs responsible for maintaining the basal conformation, and thereby, leads to efficient βarr activation. The conserved nature of this lock-and-key mechanism is further illustrated by a comprehensive sequence analysis of the human GPCRome, and demonstrated in cellular context with targeted mutagenesis including “loss-of-function” and “gain-of-function” experiments with respect to βarr activation measured by an intrabody-based conformational sensor. Taken together, our findings uncover previously lacking structural insights, which explain the ability of distinct GPCRs to activate βarrs through a common mechanism, and a key missing link in the conceptual framework of GPCR-βarr interaction and resulting functional outcomes.

## Introduction

G protein-coupled receptors (GPCRs) are characterized by their universal seven transmembrane architecture, and agonist-induced coupling to heterotrimeric G-proteins and β-arrestins (βarrs). Of these, βarrs are multifunctional cytosolic proteins critically involved in regulating the signaling and trafficking of GPCRs (Kang et al., 2014; Lefkowitz and Shenoy, 2005; Shenoy and Lefkowitz, 2011). Their interaction with GPCRs involves a major contribution from receptor phosphorylation, which not only drives the affinity of receptor-βarr interaction but also imparts functionally competent active conformation in βarrs driving downstream functional outcomes (Maharana et al., 2022; Ranjan et al., 2017; Reiter et al., 2012). Additional interaction of βarrs with the receptor transmembrane core and membrane lipid bilayer induces further structural changes (Eichel et al., 2018; Eichel et al., 2016; Shiraishi et al., 2021), which also fine-tune the functional capabilities of βarrs and possibly spatio-temporal aspects of their regulatory mechanisms (Asher et al., 2022; John Janetzko, 2022). Despite a poorly conserved primary sequence of GPCRs in terms of the number and spatial positioning of the putative phosphorylation sites, the near-universal nature of βarr interaction, ensuing signaling and regulatory responses, remain to be fully understood at molecular level (Chen and Tesmer, 2022; Seyedabadi et al., 2021). Moreover, differential receptor phosphorylation by different subtypes of GPCR kinases (GRKs), and possibly other kinases, in cell-type specific manner adds further complexity to GPCR-βarr binding modalities and context-specific functional specialization (Gurevich and Gurevich, 2020; Kawakami et al., 2022; Matthees et al., 2021).

There are two isoforms of βarrs namely βarr1 and 2, also known as Arrestin 2 and 3, respectively, and despite a high degree of sequence and structural similarity, they often exhibit a significant diversity in their functional contribution towards GPCR signaling and regulation (Drube et al., 2022; Ghosh et al., 2019; Srivastava et al., 2015). There are only a very few structures of βarr1 in active conformation, either in complex with receptor-derived phosphopeptides (He et al., 2021; Shukla et al., 2013), or, in complex with agonist-bound, phosphorylated GPCRs (Cao; Huang et al., 2020; Lee et al., 2020; Staus et al., 2020; Yin et al., 2019). The structural coverage for βarr2 is even more sparse with the structural snapshots limited to either a complex with CXCR7-derived phosphopeptide (Min et al., 2020) or in IP6-bound state (Chen et al., 2017). While these structures provide useful information about βarrs’ interaction with the receptors, there is only a limited information available about the phosphorylation patterns either due to chimeric constructs used in these studies or the lack of visualization of multiple phosphorylation sites resulting from insufficient structural resolution. Thus, the quest to decipher precise molecular details of βarr activation, especially a potentially conserved mechanism by which they can interact with a large repertoire of receptors harboring different phosphorylation patterns, remains open. This represents a major knowledge gap in our current understanding of GPCR signaling and regulatory paradigms governing and fine-tuning the signal-transduction through this versatile class of receptors.

In this backdrop, here we present cryogenic-electron microscopy (cryo-EM) structures of full length βarr1 and 2 activated by defined phosphorylation patterns encoded in the form of phosphopeptides, which are derived from three different GPCRs namely the complement C5a receptor subtype 1 (C5aR1), the CXC chemokine receptor subtype 4 (CXCR4) and the vasopressin receptor subtype 2 (V2R). These structural snapshots reveal that a P-X-P-P pattern of phosphorylation in GPCRs engages a K-K-R-R-K-K sequence in the N-domain of βarrs forming a “lock-and-key” type interaction interface that directs βarr activation. Interestingly, a large repertoire of GPCRs encode putative P-X-P-P motif either in their carboxyl-terminus, or, the 3^rd^ intracellular loop (ICL3), suggesting the conserved nature of this activation mechanism. We validate this mechanism in cellular context on a set of distinct receptors using conformational biosensors and structure-guided modification of the P-X-P-P key leading to “gain-of-function” and “loss-of-function” with respect to βarr activation. Collectively, our data reveal conserved principles of βarr activation, and provide a structural mechanism that guides phosphorylation-dependent βarr activation by GPCRs.

## Results

Although cryo-EM has been used to determine the structures of GPCR-βarr1 complexes (Cao; Huang et al., 2020; Lee et al., 2020; Staus et al., 2020; Yin et al., 2019), all the previous structures of βarrs in basal state (Han et al., 2001; Zhan et al., 2011), bound to phosphopeptides (He et al., 2021; Shukla et al., 2013) or IP6 (Chen et al., 2017) have been determined using X-ray crystallography. Therefore, in order to test the feasibility of structure determination of βarrs in complex with different phosphorylation patterns encoded in GPCR phosphopeptides by cryo-EM, we first reconstituted V2Rpp-βarr1 complex together with a set of conformationally selective Fabs, which recognize activated βarr1 (Ghosh et al., 2017). Subsequently, we analyzed these complexes using negative-staining single particle EM, which revealed monodisperse particle distribution and 2D class averages where the densities of βarr1 and Fabs were clearly discernible (Figure 1B and Figure S1A). Based on these observations, we synthesized and characterized a set of phosphopeptides corresponding to the carboxyl-terminus of the human complement C5a receptor subtype 1 (C5aR1) and the CXC chemokine receptor subtype 4 (CXCR4), and assessed their ability to activate βarr1 measured in terms of Fab30 reactivity (Figure S1B-S1E). We identified the phosphorylation patterns from C5aR1 (C5aR1pp2; referred to as C5aR1pp hereon) and CXCR4 (CXCR4pp2; referred to as CXCR4pp hereon), which elicited maximal Fab30 reactivity as a measure of βarr1 activation (Figure S1B-S1E). Subsequently, we reconstituted C5aR1pp-βarr1-Fab30 and CXCR4pp-βarr1-Fab30 complexes, validated their monodispersity and architecture using negative-staining EM (Figure 1C and 1D and Figure S1F and S1G), and subjected these complexes to cryo-EM. We successfully determined the structures of C5aR1pp-βarr1-Fab30 and CXCR4pp-βarr1-Fab30 complexes at global resolutions of 3.4 Å and 4.8 Å, respectively (Figure 1E and 1F, Figure S2).

**Figure 1.**
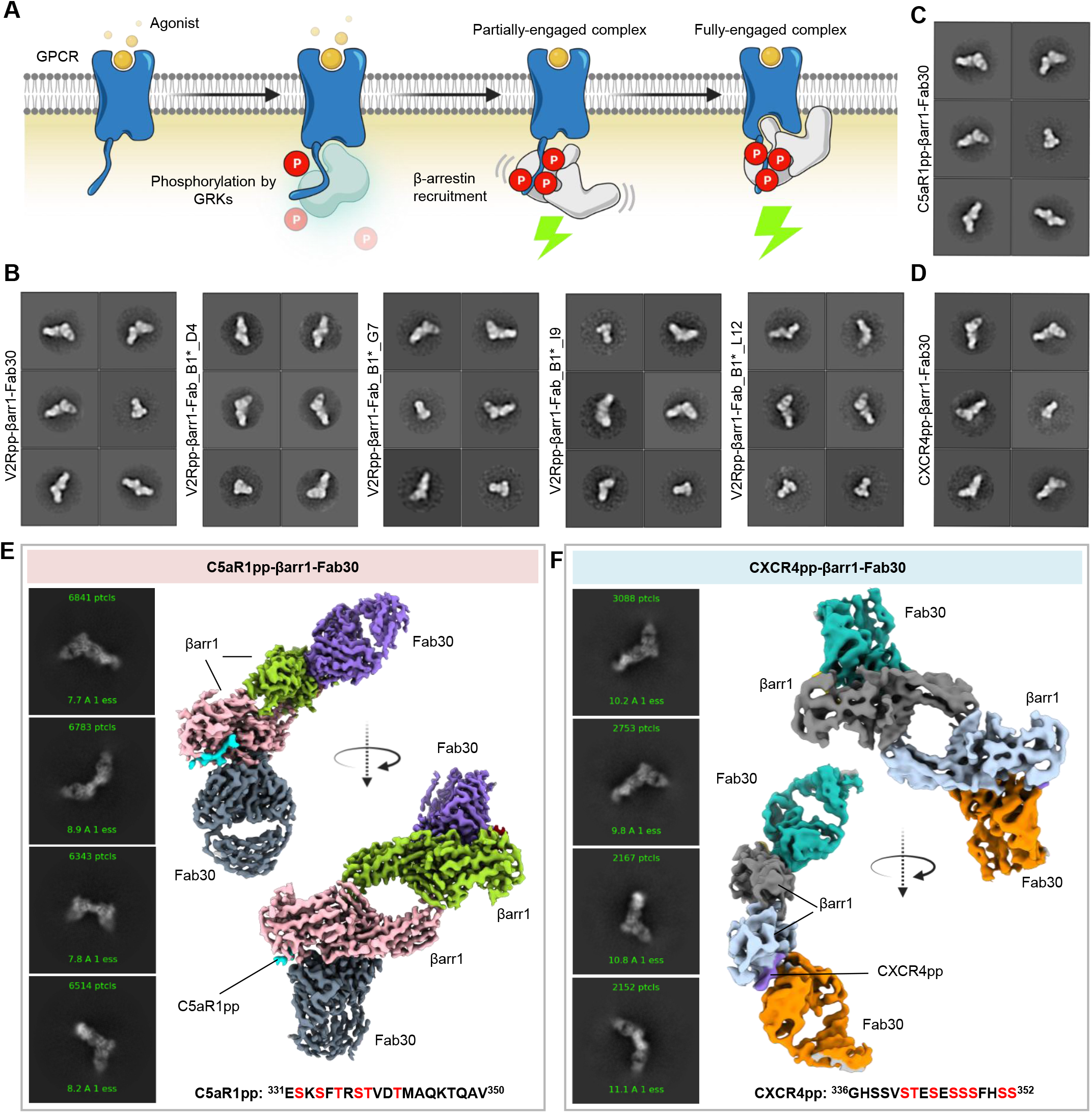
Reconstitution and structure determination of C5aR1pp/CXCR4pp-βarr1-Fab30 complexes. **(A)** Agonist-stimulation of GPCRs leads to receptor phosphorylation by GPCR kinases (GRKs) followed by the recruitment and activation of βarrs governed through the phosphorylated residues and activated receptor core. **(B)** Negative-staining EM-based 2D class averages of V2Rpp-βarr1 complexes stabilized by Fab30, Fab_B1*_D4, Fab_B1*_G7, Fab_B1*_I9 and Fab_B1*_L12 respectively. **(C-D)** Negative-staining EM-based 2D class averages of C5aR1pp-βarr1-Fab30 and CXCR4pp-βarr1-Fab30 complexes, respectively. **(E-F)** Selected 2D class averages and surface representation of C5aR1pp-βarr1-Fab30 and CXCR4pp-βarr1-Fab30 structures, respectively, determined by cryo-EM.

### βarr1 structures in complex with C5aR1 and CXCR4 phosphorylation patterns

Both structures revealed a dimeric arrangement with the two βarr1 protomers making dimeric contacts through the C-edge loops and finger loops (Figure 2A and 2B). βarr1 protomers exhibit nearly-identical overall structures with each other (RMSD, 0.08 Å for C5aR1pp-βarr1 and RMSD 0.15 Å for CXCR4pp-βarr1) with clear densities of the phosphopeptides visible in the EM map (Figure S3A and 3B). C5aR1pp and CXCR4pp are positioned in a positively-charged groove on the N-domain of βarr1 (Figure 2C and 2D), and the phosphate moieties make extensive contacts with Arg/Lys residues at the binding interface (Figure 2E and 2F). Interestingly, three phosphate groups arranged in a P-X-P-P type pattern, where P is a phosphorylated residue and X is another amino acid, in both the phosphopeptides engage a nearly-identical set of Lys/Arg residues in βarr1 (Figure 2E and 2F). Specifically, pT^336^-R^337^-pS^338^-pT^339^ pattern in C5aR1pp and pS^344^-E^345^-pS^346^-pS^347^ pattern in CXCR4pp engages K^294^-K^11^-R^25^-R^7^-K^10^-K^107^ in βarr1. The other phosphate groups present in the phosphopeptides are either not involved in a direct contact with Arg/Lys, sparsely linked with Arg/Lys, or, positioned outside the binding groove. The N- and C-domains of βarr1 exhibit an inter-domain rotation of approximately 20° when compared to the basal state of βarr1, in both the structures, which is a hallmark of βarr activation upon binding of phosphorylated GPCRs (Shukla et al., 2013) (Figure 2G). Moreover, the three major loops in βarr1 namely the finger, middle and lariat loop also exhibit significant reorientation upon binding of C5aR1pp and CXCR4pp compared to the basal state although their positioning is almost identical between the two structures (Figure 2H). Finally, the three-element interaction and polar-core network in βarr1 are also disrupted upon binding to C5aR1pp and CXCR4pp when compared to the basal state structure through the displacement of carboxyl-terminus of βarr1 from the N-domain and repositioning of the lariat loop through the interaction of K^294^ with a phosphate moiety (Figure S4A and S4B). These structural features and interaction interface are analogous to that observed in V2Rpp-βarr1 - Fab30 crystal structure determined previously (Shukla et al., 2013), although the primary sequence and phosphorylation patterns encoded by C5aR1pp and CXCR4pp are distinct from V2Rpp (Figure S4C).

**Figure 2.**
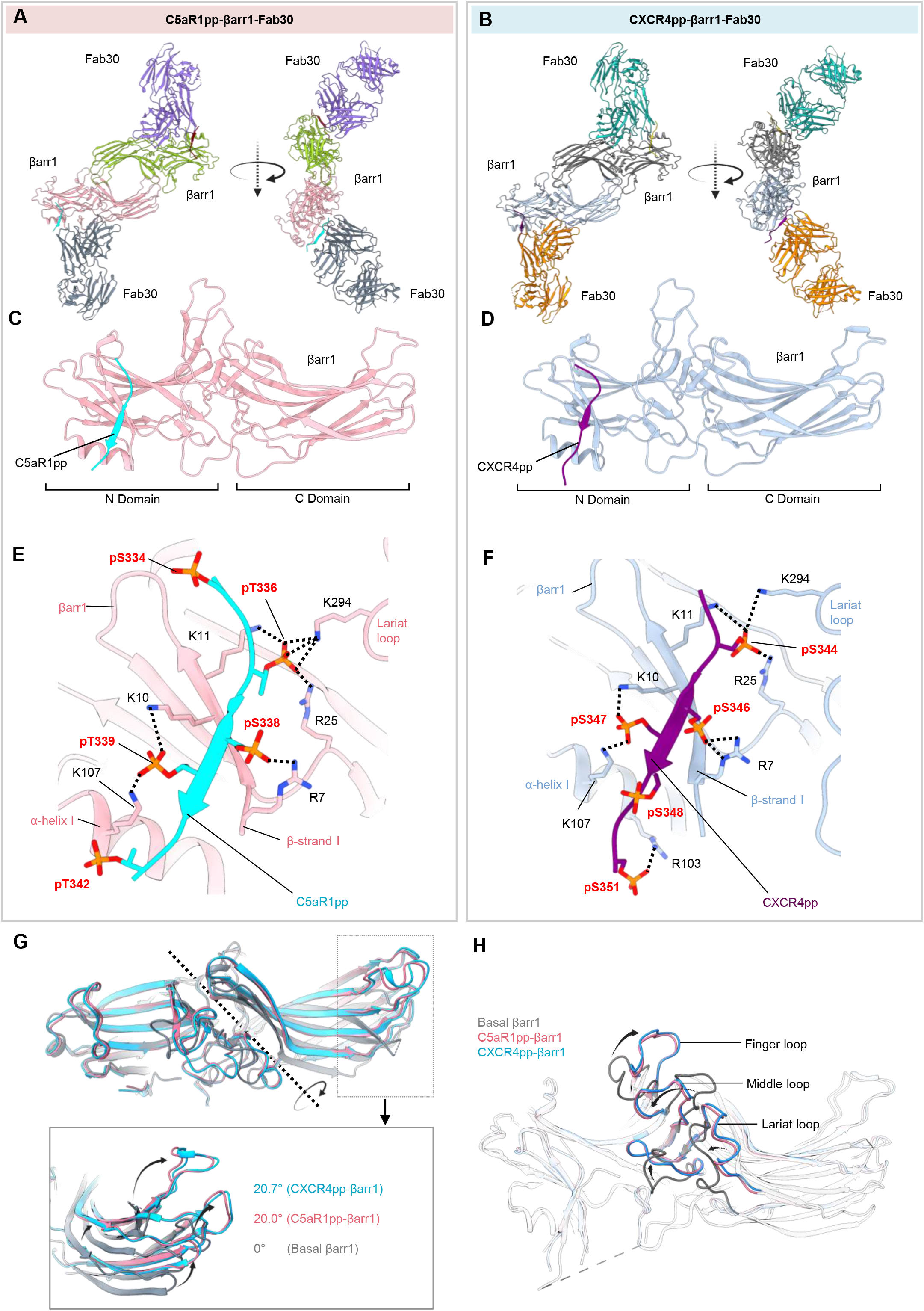
Overall structures and key structural features of C5aR1pp/CXCR4pp-βarr1-Fab30 complexes. **(A-B)** Overall structure of C5aR1pp-βarr1-Fab30 and CXCR4pp-βarr1-Fab30 complexes shown with ribbon representation. **(C-D)** Structure of individual C5aR1pp-βarr1-Fab30 and CXCR4pp-βarr1-Fab30 complex protomers shown as ribbon representation to indicate the binding of phosphopeptides on the N-domain of βarr1. **(E-F)** Stabilizing charge-charge interactions of C5aR1pp and CXCR4pp with the N-domain groove residues of βarr1 indicated as dotted lines. pS and pT refers to phospho-Ser and phospho-Thr residues, respectively. **(G)** Inter-domain rotation in βarr1 upon binding of C5aR1pp (pink) and CXCR4pp (blue) is compared with the basal conformation of βarr1 determined previously (PDB 1G4M, gray). **(H)** Superimposition of C5aR1pp- and CXCR4pp-bound βarr1 structures with the basal conformation of βarr1 (PDB 1G4M, gray) indicating the repositioning of finger, middle and lariat loops upon βarr1 activation.

### Structure-guided engineering yields structures of activated βarr2

As mentioned earlier, the structural coverage of active βarrs, either in complex with full GPCRs or GPCR-derived phosphopeptides, is limited primarily to βarr1. Activated structures of the other isoform i.e., βarr2 are represented only by an IP6-bound crystal structure (Chen et al., 2017) and a complex of truncated βarr2 with a phosphopeptide derived from a βarr-biased 7TMR (CXCR7) (Min et al., 2020). Therefore, we set out to reconstitute and determine the structure of βarr2 in complex with the phosphopeptides derived from different receptors, i.e., C5aR1pp and CXCR4pp using cryo-EM. Surprisingly however, we did not observe a significant Fab30 reactivity to C5aR1pp/CXCR4pp-bound βarr2 while it robustly recognized V2Rpp-βarr2 complex (Figure 3A). Therefore, we analyzed the Fab30 interaction interface on C5aR1pp-bound βarr1 structure to identify a potential reason for the lack of Fab30 reactivity with βarr2. Interestingly, we observed that Fab30 epitope is conserved between βarr1 and 2 except two residues i.e., instead of F^277^ and A^279^ as in βarr1, βarr2 contains L^278^ and S^280^ in the corresponding positions, respectively (Figure 3B). Therefore, we generated a βarr2 mutant, referred to as βarr2^DM^, and tested its reactivity to Fab30 upon activation by distinct phosphopeptides. In line with our hypothesis, we observed a robust interaction of Fab30 with C5aR1pp-βarr2^DM^ and CXCR4pp-βarr2^DM^ complex, and we also noticed that the interaction of Fab30 with V2Rpp-βarr2^DM^ was further enhanced compared to βarr2^WT^ (Figure 3C). We also confirmed that βarr2^DM^ exhibits a similar pattern of interaction as βarr2^WT^ with V2R and C5aR1 in cellular context (Figure 3D) and shows near-identical endosomal trafficking as βarr2^WT^ upon stimulation of V2R and C5aR1 (Figure 3E). Thus, βarr2^DM^ provides us an excellent handle to reconstitute stable complexes with receptor phosphopeptides suitable for cryo-EM. In fact, we successfully managed to reconstitute monodisperse V2Rpp-βarr2^DM^-Fab30 and C5aR1pp-βarr2^DM^-Fab30 complexes (Figure S5C and S5I) and determine their cryo-EM structures at 4.2 Å and 4.4 Å resolution, respectively (Figure 4A and B, Figure S5D-S5H, S5J-S5N). In order to simplify the discussion, we refer to βarr2^DM^ as βarr2 from here onwards unless specified otherwise.

**Figure 3.**
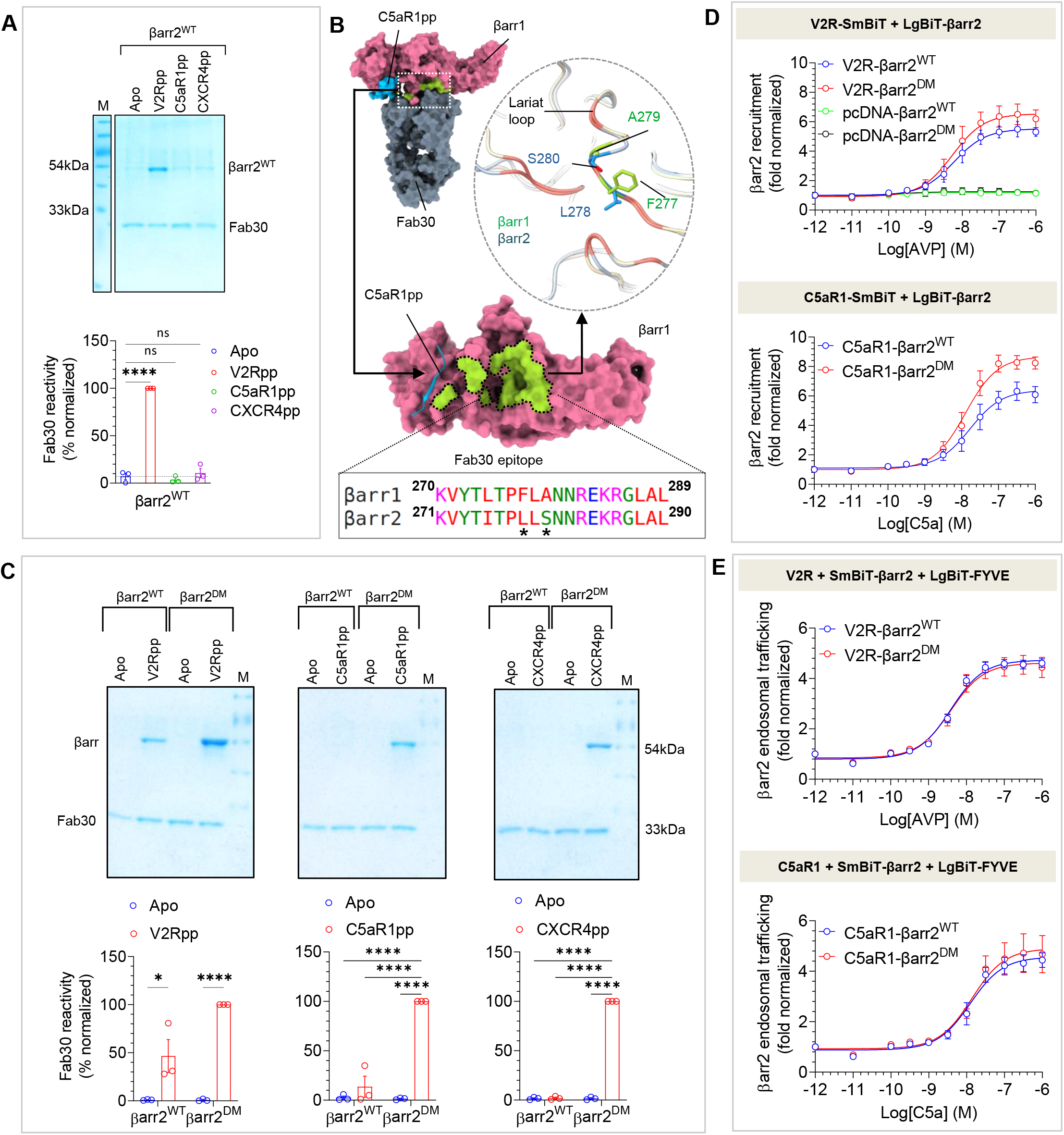
Generation and characterization of βarr2DM for V2Rpp/C5aR1pp bound complexes. **(A)** Fab30 reactivity of C5aR1pp and CXCR4pp activated βarr2WT was measured by co-immunoprecipitation (co-IP) assay. C5aR1pp and CXCR4pp activated βarr2WT was not recognized by Fab30 (upper panel). Densitometry-based quantification of the co-IP data is presented in the lower panel (mean ± SEM; n=3; normalized with respect to V2Rpp signal as 100%; One-way ANOVA, Dunnett’s multiple comparisons test; (****p<0.0001; ns, non-significant). **(B)** Comparison of the epitope region of Fab30 in βarr1 with βarr2 reveals that instead of F277 and A279 as in βarr1, βarr2 contains L278 and S280 in corresponding positions (indicated with asterisk). **(C)** Co-IP assay showing the reactivity of Fab30 towards activated βarr2DM upon binding of C5aR1pp and CXCR4pp (upper panel). Densitometry-based quantification is presented in the lower panel (mean ± SEM; n=3; normalized with respect to Fab30 reactivity towards activated βarr2DM treated as 100%; Two-way ANOVA, Tukey’s multiple comparisons test; **p<0.01, ***p<0.001, ****p <0.0001, ns = non-significant). **(D)** A side-by-side comparison of agonist-induced βarr2WT and βarr2DM recruitment to V2R (left panel) and C5aR1(right panel) in the nanoBiT assay (Receptor-SmBiT+LgBiT-βarr2WT/βarr2DM) (mean±SEM; n=5; normalized as fold over basal). **(E)** A side-by-side comparison of βarr2WT and βarr2DM endosomal trafficking in response to agonist (AVP for V2R and C5a for C5aR1) in the nanoBiT assay (Receptor+SmBiT-βarr2+LgBiT-FYVE) (mean±SEM; n=5; normalized as fold over basal).

**Figure 4.**
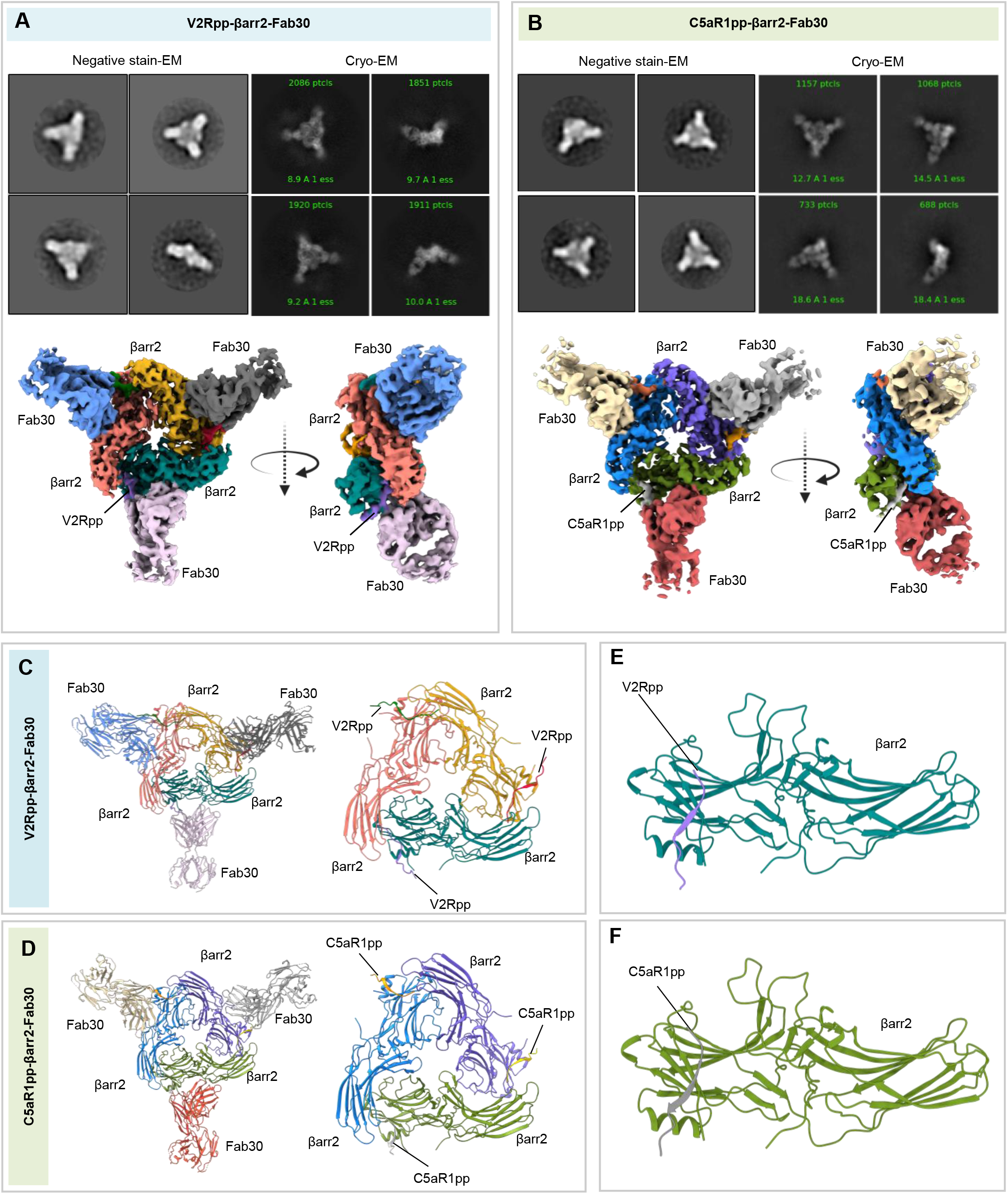
Overall structures of V2Rpp/C5aR1pp-βarr2 complexes. **(A-B)** Overall cryo-EM structure of V2Rpp-βarr2-Fab30 (top) and C5aR1pp-βarr2-Fab30 complexes (bottom), respectively, in a trimeric assembly with βarr2 and Fab30 molecules colored as individual units. Front and side views of the trimeric complex EM map have been shown with βarr2 molecules in blue, olive green and purple; and Fab3O molecules in beige, red and gray. **(C-D)** Overall trimeric arrangement of V2Rpp/C5aR1pp-βarr2-Fab30 complexes in the cryo-EM structures shown here as cartoon representation (left panel). Domain organization of the βarr2 molecules in trimeric assembly without Fab30 shown as cartoon representation (right panel). **(E-F)** Structure of individual βarr2 protomers in V2Rpp/C5aR1pp-βarr2-Fab30 complexes showing the binding of phosphopeptides on the N-domain of βarr2

### Structures of βarr2 in complex with V2Rpp and C5aR1pp

The V2Rpp-βarr2 and C5aR1pp-βarr2 structures exhibited a trimeric assembly of βarr2 with the individual protomers arranged through N- to C-domain contacts (Figures 4A-4D and Figure S7). The overall structural features of the individual protomers in each structure were nearly identical as reflected by low RMSD (0.005 Å for V2Rpp-βarr2 and 0.038 Å for C5aR1pp-βarr2) with the phosphopeptide densities clearly visible in the EM maps (Figures S3C and S3D). Like βarr1 structures, V2Rpp and C5aR1pp are positioned in a positively-charged groove on the N-domain of βarr2 (Figures 4E and 4F), and the phosphate moieties make extensive contacts with Arg/Lys residues at the binding interface (Figures 5A and 5B). Remarkably, we observed that three phosphate groups arranged in a P-X-P-P pattern in the phosphopeptides, engaged an analogous set of Lys/Arg residues as in βarr1 (Figures5A and 5B). Specifically, pT^360^-A^361^-pS^362^-pS^363^ in V2Rpp and pT^336^-R^337^-pS^338^-pT^339^ in C5aR1pp engage K^295^-K^12^-R^26^-R^8^-K^11^-K^108^ in βarr2 (Figures 5A and 5B). Similar to βarr1 structures, the other phosphate groups present in the phosphopeptides are either not involved in a direct contact with Arg/Lys or positioned outside the binding groove. The N- and C-domains of βarr2 exhibit an inter-domain rotation of approximately 25° when compared to the basal state of βarr2 in both the structures, which is relatively higher from that observed in phosphopeptide bound βarr1 (Figure 5C). Moreover, the three major loops in βarr2 namely the finger, middle and lariat loop also exhibit significant reorientation upon binding of V2Rpp and C5aR1pp compared to the basal state although their positioning is almost identical between the two structures (Figure 5D). Finally, the three-element interaction and polar-core network in βarr2 are also disrupted upon binding to V2Rpp and C5aR1pp when compared to the basal state structure through the displacement of carboxyl-terminus of βarr2 from the N-domain and repositioning of the lariat loop through the interaction of K^295^ with a phosphate moiety (Figures 5E and 5F).

**Figure 5.**
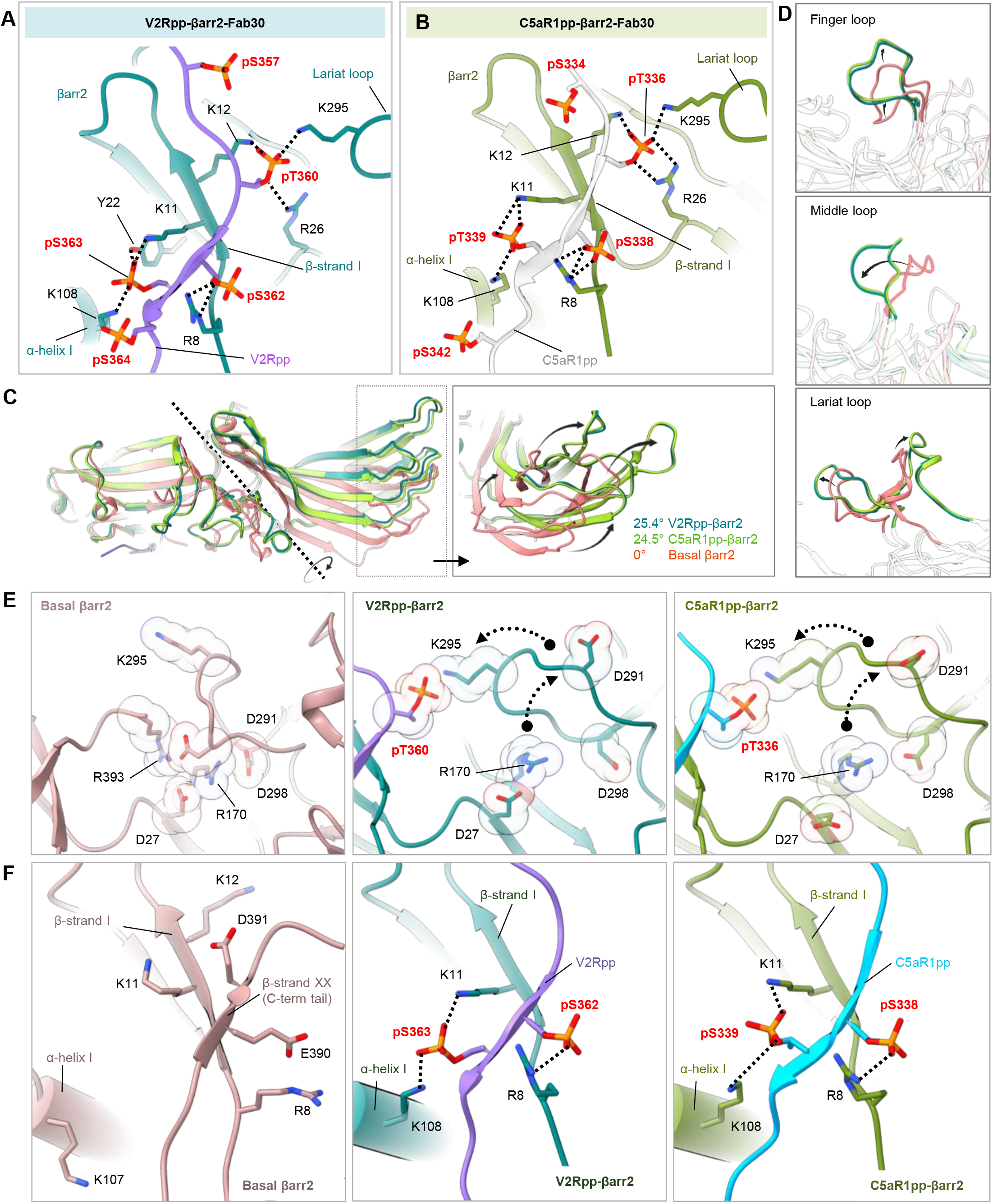
Overall structure and key structural features of V2Rpp/C5aR1pp-βarr2-Fab30 complexes. **(A-B)** Extensive charge-charge interactions between the phosphate residues in V2Rpp/C5aR1pp with Lys/Arg in the N-domain (represented as black dotted lines) stabilize the V2Rpp and C5aR1pp into the N-domain groove of βarr2**. (C)** V2Rpp (dark green) and C5aR1pp (light green) activated βarr2 structures were superimposed with the basal state of βarr2 (PDB 3P2D, orange) and inter-domain rotations were calculated. **(D)** Conformational changes observed in the finger (top), middle (middle) and lariat loops (bottom) in the activated βarr2 compared to the basal state crystal structure of βarr2. **(E)** Polar core environment in basal βarr2 (PDB 3P2D, left) and disruption of polar core interactions upon binding of V2Rpp (middle) and C5aR1pp to βarr2 (right). **(F)** Three-element interaction network consisting of βarr2 C-terminal β-strand XX, α-helix1 and β-strand1 in the basal state of βarr2 (left). Binding of the phosphopeptides V2Rpp and C5aR1pp to βarr2 results in the displacement of the β-strand XX, and engages the phosphopeptide V2Rpp (middle) and C5aR1pp (right) into the N-domain groove of βarr2 through hydrogen bonds and polar interactions.

### A conserved “lock-and-key” mechanism of βarr activation

As mentioned earlier, the analysis of these structural snapshots in terms of phosphorylation sites revealed a conserved P-X-P-P pattern with nearly-identical interactions with analogous residues in βarr1 and 2 (Figures 6A-6C). Therefore, we analyzed the primary sequence of all non-olfactory and non-orphan GPCRs in their carboxyl-terminus and intracellular loop 3 (ICL3) to identify the occurrence of P-X-P-P pattern in these receptors (Supplementary Table S2). We observed that a large set of GPCRs harbored this motif in their carboxyl-terminus sequence and several receptors also included it in their ICL3 (Figures 6F and 6G). In order to validate the functional contribution of this motif in GPCR-induced βarr activation, we employed Ib30-based conformational biosensor in cellular context using three different receptors, which possess P-X-P-P motif either in their C-terminus (CXCR3), ICL3 (M2R), or, lack it (CXCR7). At the level of βarr1 conformation, Ib30 sensor reports the degree (>15°) of inter-domain rotation as a proxy of βarr activation upon its interaction with activated and phosphorylated receptors (Dwivedi-Agnihotri et al., 2020). In agreement with our hypothesis, we observed robust reactivity of Ib30 sensor with βarr1 for CXCR3 and M2R but not for CXCR7 (Figure 7A). To further corroborate these findings, we used two different receptors namely, the Bradykinin receptor subtype 2 (B2R) and C5aR1 for structure-based targeted mutagenesis to probe “gain-of-function” and “loss-of-function”, respectively, in terms of Ib30 reactivity pattern. As presented in Figures 7B and 7C, activation of the wild-type B2R fails to induce an interaction between βarr1 and Ib30 as it lacks a P-X-P-P motif in its C-terminus, although B2R is capable of recruiting βarrs (Baidya et al., 2020a; Zimmerman et al., 2011). Interestingly however, reconstitution of P-X-P-P motif in B2R by double mutation (ΔG^368^+L^370^T) results in a robust Ib30 reactivity upon agonist-stimulation (Figure 7B). Along the same lines, a mutant version of C5aR1pp, where the P-X-P-P motif is disrupted by insertion of an additional arginine residue between pT^336^ and pS^338^, completely loses the ability to induce a conformation that is recognizable by Fab30 (Figure 7C). Moreover, the corresponding mutation in C5aR1 also leads to a dramatic loss of Ib30 reactivity in cellular context (Figure 7C). Taken together these data establish the P-X-P-P phosphorylation pattern in GPCRs as a “key” to open the “lock” in βarrs leading to its conformational activation (Figure 7D).

**Figure 6.**
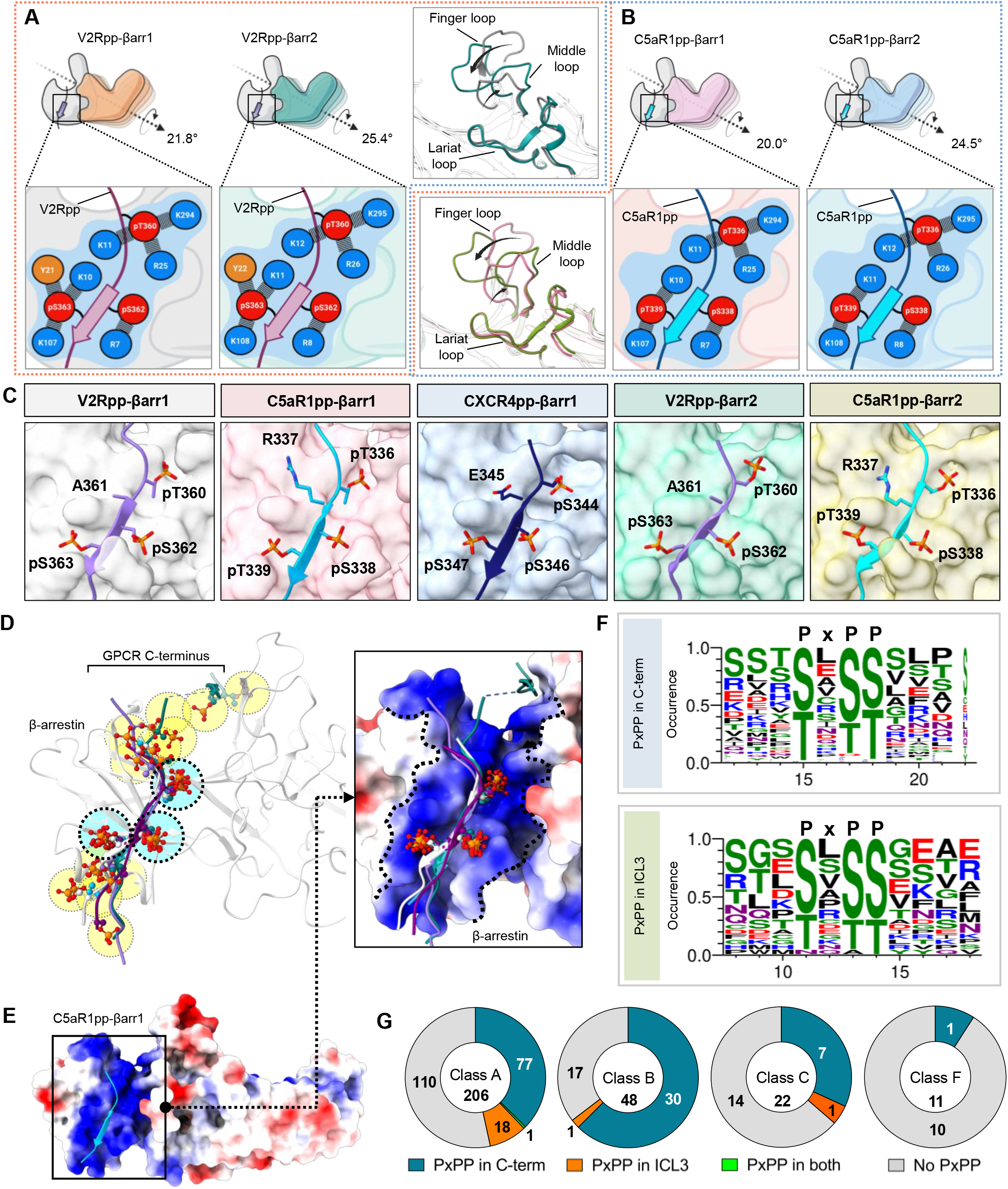
A conserved lock and key mechanism of βarr activation. **(A)** Comparison of the V2Rpp-bound βarr1 and 2 reveals similar interactions of V2Rpp with both isoforms of βarrs although a slightly higher inter-domain rotation is observed in βarr2 (left). A schematic representation of the interface network between negatively charged phospho-residues (red) and positively charged residues (blue) of βarrs are shown (below, zoomed in box). Although the lariat loops of the two structures align well, significant deviations can be observed for the finger and middle loops (right, inset box). **(B)** Comparative analysis of C5aR1pp-bound βarr1 and 2 structures reveal similar interactions of C5aR1pp with both βarr isoforms, but again, a higher inter-domain rotation is observed for βarr2. A similar representation of the interface between negatively charged phospho-residues (red) and positively charged residues (blue) of βarrs are shown (below, zoomed in box). **(C)** In all the structures of phosphopeptide-bound βarrs, a conserved motif can be observed with respect to the three phospho-residues (dotted yellow circles), referred to as P-X-P-P motif, where “P” is a phospho-Ser/Thr and “X” can be any residue. **(D)** Superposition of V2Rpp-βarr1 (PDB 4JQI), V2Rpp-βarr2 (PDB 8GOC), C5aR1pp-βarr1 (PDB 8GO8), C5aR1pp-βarr2 (PDB 8GOO) with D6Rpp-βarr2 (PDB 8GO9) clearly shows conservation of phosphates corresponding to P-X-P-P position where as other phosphates are distributed throughout the phosphopeptides. **(E)** Superposition of phosphopeptides on C5aR1pp-βarr1 reveals the conserved phospho-residues on positively charged cleft present on βarrs’ N-Domain. βarr is shown as coulombic charged surface here. **(F)** A sequence alignment of the C-terminal tail and ICL3 residues of non-olfactory and non-orphan Class-A receptors reveal the consensus sequence, “P-X-P-P” required for activation of βarrs. The consensus sequence logo was generated with the WEBLOGO tool52 and sequence alignment was performed with Kalign (Lassmann, 2020). A stretch of 11 amino acid residues have been shown for better representation. **(G)** Proportions of GPCRs of Class A, B, C and F having P-X-P-P motif in C-terminus or ICL3 have been represented as pie charts.

**Figure 7.**
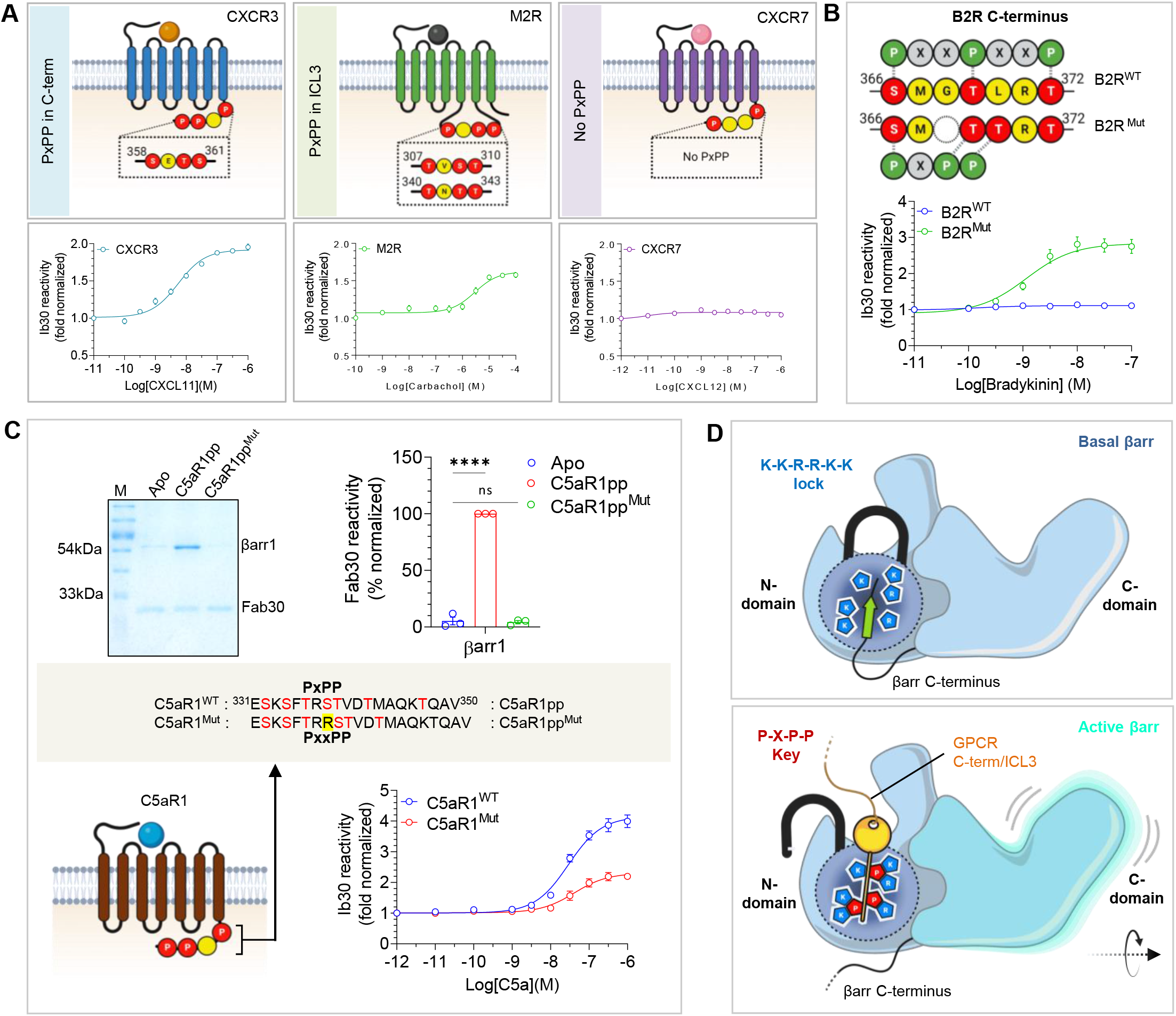
The P-X-P-P motif in GPCRs is sufficient for βarr activation. **(A)** NanoBiT-based assay for assessing Ib30 reactivity to CXCR3 (left panel), M2R (middle panel), and CXCR7 (right panel) activated βarr1 (Receptor+SmBiT-βarr1+LgBiT-Ib30) (mean±SEM; n=3; normalized as fold over basal). **(B)** Deletion of G368 and substitution of L370 to Ala in B2R engineers the “P-X-P-P key” and results in “gain-of-function” in terms of Ib30 reactivity as measured using the NanoBiT assay (Receptor+SmBiT-βarr1+LgBiT-Ib30) (mean±SEM; n=3; normalized as fold over basal). **(C)** Addition of an extra Arg between positions 336 and 337 in C5aR1pp to disrupt the “P-X-P-P key” (referred to as C5aR1ppMut) leads to a near-complete loss of Fab30 (top) reactivity as measured in co-IP assay (mean±SEM; n=3; densitometry-based data normalized with respect to C5aR1pp signal as 100%; One-way ANOVA, Dunnett’s multiple comparisons test; ****p < 0.0001, ns = non-significant). Corresponding mutation in in C5aR1 to disrupt the “P-X-P-P key” (referred to as C5aR1Mut) results in a dramatic decrease in Ib30 reactivity (bottom) as measured using the NanoBiT assay (mean±SEM; n=2; normalized as fold over basal). **(D)** Schematic representation of the “lock and key” mechanism of βarr activation. The C-terminus of βarr is positioned on the N-domain, which stabilizes the basal conformation through the three-element interaction and polar-core network. Binding of GPCRs/ACRs harboring the “P-X-P-P key” engages the critical points in the “K-K-R-K-R-K lock” leading to βarr activation and functional responses.

## Discussion

The identification of conserved principles that guide the interaction and conformational activation of βarrs with a large repertoire of GPCRs has been a major challenge, and the structural snapshots of βarr1 and 2 presented here with distinct phosphorylation patterns derived from different receptors provide a breakthrough. The “P-X-P-P key” present in these phosphorylation patterns appear to be sufficient to simultaneously engage the crucial “lock points” in βarrs to facilitate their activation. It is intriguing that the binding interface of “P-X-P-P key” is conserved not only across different peptides irrespective of their primary sequence but also for both βarr isoforms (Figures 6D and 6E). This conserved binding interface and corresponding interactions ensure the displacement of βarrs’ C-terminus from the N-domain and repositioning of the lariat loop, leading to the release of the two major “breaks” on βarr activation namely, the three-element interaction and the polar-core network (Gurevich and Gurevich, 2020) (Figure S6A). It is important to note that GPCRs lacking the P-X-P-P pattern are also capable of recruiting βarrs in functionally competent conformation; however, they are likely to induce an active βarr conformation that is distinct from the receptors harboring the P-X-P-P key. Additionally, we cannot rule out the possibility that for some receptors, a functional “P-X-P-P key” may not be formed despite having a suitable primary sequence because all the phosphorylatable residues may not undergo phosphorylation in cellular context. A previous study has proposed a framework of “full” and “partial” phosphorylation codes imparting distinct βarr recruitment patterns for GPCRs (Zhou et al., 2017). Our study now identifies and establishes a general principle of efficient βarr activation through a specific phosphorylation pattern encoded in GPCRs engaging a conserved interface on βarrs.

An intriguing observation in these structural snapshots is novel dimer and trimer assemblies of βarrs. Although βarrs have strong propensity to adopt different oligomeric states (Chen et al., 2014; Chen et al., 2021; Gurevich and Gurevich, 2022), the dimer and trimer interfaces observed here differ significantly from previously reported interfaces (Chen et al., 2017; Chen et al., 2014; Chen et al., 2021) (Figure S7). The overall buried surface area in dimer and trimer assemblies are approximately 1500 Å^2^ and 4500 Å^2^, respectively, suggesting a robust and stable oligomeric arrangement. The two protomers in C5aR1pp- and CXCR4pp-bound βarr1 interface with each other through multiple hydrogen bonds, salt-bridges, and non-bonded contacts in a manner where the C-edge loop residues of one protomer are positioned into the central crest of the other protomer in proximity of the finger loop. An analogous set of interactions are also involved in trimer arrangement of βarr2 in complex with V2Rpp and C5aR1pp. Interestingly, a previous crystal structure of βarr2 in complex with IP6 also shows a trimeric arrangement although the trimer interface is different from that observed here in phosphopeptide-bound conformations (Chen et al., 2017). A comprehensive map of dimer and trimer interface with residue-level contacts is listed in Supplementary Table S3. Considering the functional multiplicity of βarrs in terms of distinct signaling and regulatory outcomes and receptor-specific responses (Gurevich and Gurevich, 2019), it is plausible that distinct oligomeric interfaces in βarrs may be a modular mechanism to fine-tune the functional contributions by providing distinct possibilities for adaptable protein-protein interaction interfaces for binding partners.

The comparison of V2Rpp- and C5aR1pp-bound βarr1 and 2 reveal a significantly higher inter-domain rotation in βarr2 compared to βarr1 as hypothesized earlier based on cellular and biochemical studies (Ghosh et al., 2019), and this may provide a plausible explanation for functional differences between the βarr isoforms as observed for multiple GPCRs. It is also noteworthy that the structural snapshots presented here involve isolated phosphopeptides with defined phosphorylation patterns without the transmembrane core of the receptors. As the interaction of receptor core imparts additional conformational changes in βarrs (Ghosh et al., 2019; Latorraca et al., 2018; Shiraishi et al., 2021), it is plausible that the full complexes of receptors and βarrs may exhibit additional conformational changes in βarrs, especially in terms of the positioning of the proximal region of the phosphorylated segment. However, the conserved principle of “P-X-P-P key” to open the “K-K-R-R-K-K lock” is likely to be maintained and guide βarr activation even in the context of full receptors.

In summary, we identify and experimentally validate a conserved principle of phosphorylation-induced βarr activation based on structural snapshots of activated βarrs in complex with distinct receptor phosphorylation patterns. Our study addresses a long-standing question in the field to decipher the molecular basis of universal βarr activation by receptor phosphorylation, and lays the foundation to further refine the conceptual framework of βarr-mediated signaling and regulation of 7TMRs.

### Data availability statement

The three-dimensional cryo-EM density maps have been deposited in the Electron Microscopy Data Bank under the accession numbers EMD-34173 (C5aR1pp-βarr1-Fab30), EMD-34175 (V2Rpp-βarr2-Fab30), EMD-34178 (C5aR1pp-βarr2-Fab30) and EMD-34188 (CXCR4pp-βarr1-Fab30). Coordinates for the atomic models have been deposited in the RCSB Protein Data Bank with the accession numbers 8GO8 (C5aR1pp-βarr1-Fab30), 8GOC (V2Rpp-βarr2-Fab30), 8GOO (C5aR1pp-βarr2-Fab30) and 8GP3 (CXCR4pp-βarr1-Fab30). Any additional information required to reanalyze the data reported in this paper is available from the corresponding author upon reasonable request.

## Acknowledgements

Research in A.K.S.’s laboratory is supported by the Senior Fellowship of the DBT Wellcome Trust India Alliance (IA/S/20/1/504916) awarded to A.K.S., Science and Engineering Research Board (EMR/2017/003804, SPR/2020/000408, and IPA/2020/000405), Council of Scientific and Industrial Research [37(1730)/19/EMR-II], Indian Council of Medical research (F.NO.52/15/2020/BIO/BMS), Young Scientist Award from Lady Tata Memorial Trust, and IIT Kanpur. We thank A. Ranjan, M. Chaturvedi and H. Dwivedi-Agnihotri for their help with the characterization of the phosphopeptides, M. Ganguly for assisting with GPCR sequence analysis, and E. Ghosh for initial characterization of βarr2^DM^. Cryo-EM was performed at BioEM lab of the Biozentrum at the University of Basel, and we thank Carola Alampi and David Kalbermatter for their excellent technical assistance.

## Authors’ contribution

JM and MKY prepared and characterized the βarr complexes, JM performed negative-staining EM with RB, and processed the cryo-EM data with RB; PS carried out all the functional assays related to βarr2^DM^ characterization and Ib30 sensor assay; MKY purified βarrs and carried out the co-IP experiments with VS and SS; ShS contributed to functional characterization of βarr2^DM^; MC screened the samples and collected cryo-EM data; AKS supervised and managed the overall project; all authors contributed to data analysis, interpretation and manuscript writing.

## Accession number

The cryo-EM maps and structures have been deposited in the EMDB and PDB with accession numbers EMD-34173 and 8GO8 (C5aR1pp-βarr1-Fab30), EMD-34174 and 8GO9 (D6Rpp-βarr2-Fab30), EMD-34175 and 8GOC (V2Rpp-βarr2-Fab30), EMD-34178 and 8GOO (C5aR1pp-βarr2-Fab30) and EMD-34188 and 8GP3 (CXCR4pp2-βarr1-Fab30) respectively.

## Materials and Methods

### General reagents, plasmids, and cell culture

Most of the general reagents were purchased from Sigma Aldrich unless mentioned otherwise. Dulbecco’s Modified Eagle’s Medium (DMEM), Dulbecco’s Phosphate buffer saline (PBS), Fetal-Bovine Serum (FBS), Trypsin-EDTA, Hank’s balanced salt solution (HBSS), and penicillin-streptomycin solution were purchased from Thermo Fisher Scientific. HEK293 cells were obtained from ATCC and maintained in DMEM (Gibco, Cat no. 12800-017) supplemented with 10% FBS (Gibco, Cat no. 10270-106) and 100 U ml^-1^ penicillin (Gibco, Cat no. 15140122) and 100 μg ml^-1^ streptomycin (Gibco, Cat no. 15140-122) at 37 °C under 5% CO2. The cDNA coding region for the mentioned receptors, namely V2R, C5aR1 (WT and Mut), B2R (WT and Mut), M2R, CXCR3, and CXCR7 were cloned in pcDNA3.1 consist of HA signal sequence followed by FLAG tag at the N-terminus of the receptor. For the NanoBiT assay, receptors harboring SmBiT at the C-terminus were generated by subcloning in the lab, and other constructs have been described previously(Baidya et al., 2022; Baidya et al., 2020a; Baidya et al., 2020b). All the constructs were verified by DNA sequencing (Macrogen).

### Expression and purification of βarrs

Full length rat βarr1, βarr2^WT^ and βarr2^DM^ were cloned into pGEX-4T3 vector with thrombin cleavage site between GST tag and βarr. Similar protocol was followed for purifying all three forms of βarr. βarrs were expressed in *E. coli* BL21 cells and grown in Terrific broth media supplemented with 100 μg ml^-1^ Ampicillin. A primary culture of 50 ml volume was inoculated with an isolated colony from freshly transformed LB-Amp plate. Primary culture was grown till a cell optical density at 600 nm (OD_600_) of 0.8-1 and further inoculated into a secondary culture of TB-Amp of 1.5 L volume till OD_600_ 0.8-1. The expression of βarrs was then induced with 25 μM IPTG concentration and cells were allowed to grow till 16 h at 18 °C. Cultures were harvested and stored at −80 °C until further use. Harvested pellets were of 12-15 g in mass.

For purification, cells were lysed by sonication in lysis buffer; 25 mM Tris, pH 8.5, 150 mM NaCl, 1 mM PMSF (phenylmethylsulfonyl fluoride), 2 mM Benzamidine, 1 mM EDTA (Ethylenediaminetetraacetic acid), 5% Glycerol, 2 mM Dithriothreitol (DTT) and 1 mg ml^-1^ Lysozyme. The lysate was centrifuged at 18000-20000 rpm at 4 °C and supernatant was allowed to bind to Glutathione resin (GS resin) (Glutathione Sepharose^TM^ 4 Fast Flow, GE Healthcare Cat no. 17-5132-02) in a batch binding mode for overnight at 4 °C. GS-resin bound GST-βarr was transferred into Econo columns (Biorad, Cat. no.) and washed rigorously with wash buffer (25 mM Tris, pH 8.5, 150 mM NaCl, 2 mM DTT and 0.02% n-dodecyl-β-D-maltopyranoside (DDM). Afterward, on-column cleavage was set up by adding thrombin to 1: 1 resin: buffer slurry at room temperature for 2 h. βarrs were then eluted with gravity flow and further with buffer 25 mM Tris, pH 8.5, 350 mM NaCl and 0.02% DDM and 2 mM DTT. Eluted proteins were concentrated and further purified on a HiLoad 16/600 Superdex column in buffer 25 mM Tris, pH 8.5, 350 mM NaCl, 2 mM DTT and 0.02% DDM. Fractions corresponding to pure βarr were flash frozen with 10% glycerol and stored at −80 °C until further use.

### Expression and purification of Fabs

A similar protocol for expression and purification was followed for all the Fabs and they were purified as previously mentioned (Ghosh et al., 2017). Briefly, Fabs were expressed in the periplasmic fraction of *E. coli* M55244 cells (ATCC) and purified using Protein L resin (GE Healthcare Cat no. 17547802) with gravity flow affinity chromatography. Cells transformed with Fab plasmid were grown in 50 ml 2xYT media and allowed to grow overnight at 30 °C. 1 L 2xYT media was inoculated with 5% volume of initial inoculum and grown for an additional 8 h at 30 °C. Cells were collected and resuspended in an equal volume of CRAP medium supplemented with 100 μg ml^-1^ ampicillin, and grown further for 16 h at 30 °C. For purification, cells were lysed in lysis buffer (50 mM HEPES-Na^+^, pH 8.0, 0.5 M NaCl, 0.5% (v/v) Triton X-100, 0.5 mM MgCl_2_) by sonication. Cell lysate was heated in a 65 °C water bath for 30 min and cooled immediately on ice for 5 min. Lysate was centrifuged at 20,000 rpm and passed through pre-equilibrated Protein L resin packed gravity flow affinity columns. Binding was performed at room temperature and beads were washed extensively with wash buffer (50 mM HEPES-Na^+^, pH 8.0, 0.5 M NaCl). Fabs were eluted with 100 mM acetic acid into tubes containing 10% vol of 1 M HEPES, pH 8.0 for neutralization. Eluted samples were desalted into buffer (20 mM HEPES-Na^+^, pH 8.0, 0.1 M NaCl) using a pre-packed PD-10 column (GE Healthcare). Purified Fabs were flash-frozen and stored at −80 °C supplemented with 10% (v/v) glycerol until further use.

### Co-immunoprecipitation assay

For co-immunoprecipitation assay, 2.5 μg of β-arrestins were incubated with different phosphopeptides at 10-fold molar excess in binding buffers (20 mM HEPES, pH 7.4, 150 mM NaCl) for 1 h at room temperature for activation. Post peptide-induced activation, 5 μg Fab30 was added, and reaction was incubated for an additional 1 h at room temperature. After 1 h hour, 25 μl of pre-equilibrated protein L beads (Capto™ L resin, GE Healthcare Cat no. 17547802 Cat. no. 17547803) was added and reaction was incubated for 90 min at room temperature, followed by five washes with binding buffer containing 0.01% LMNG. Bound protein was eluted with 2X SDS loading buffer and 15 μl sample was analyzed on 12% SDS-PAGE. For statistical analyses, protein bands were quantified using ImageJ software suite and the values were plotted using GraphPad Prism software (v9.3). The data were normalized with respect to their respective experimental control and appropriate statistical analyses were performed as indicated in the corresponding figure legend.

### Reconstitution of pp-βarr-Fab complexes

The previously published protocol was followed for complex purification with minor modifications(Shukla et al., 2013). Briefly, βarrs were activated with corresponding phosphopeptides at a 1:3 molar ratios of βarr: phosphopeptide for 30-40 min at room temperature. Respective Fabs were added to the phosphopeptide-βarr mixture at 1:1.5 molar ratio of βarr:Fab and incubated for 1 h at room temperature. To remove excess Fabs, the phosphopeptide-βarr-Fab complexes were concentrated with 30,000 MWCO concentrators (Vivaspin, Cytiva Cat no. 28932361) and injected into Superose 6 Increase 10/300 GL (Cytiva Cat no. 29091596) gel-filtration column. Fractions were further analyzed on SDS-PAGE and selected fractions were pooled and concentrated for structural studies.

### Negative-staining EM

Complex formation, homogeneity and particle quality of the samples were judged through negative staining of the samples prior to data collection under cryo conditions for high resolution reconstructions. Negative staining of the samples was performed with uranyl formate in accordance with the previously published protocols(Peisley and Skiniotis, 2015). For imaging, 3.5 μl of the samples were dispensed on glow discharged carbon/formvar coated 300 mesh Cu grids (PELCO, Ted Pella) and allowed to adsorb for 1 min, followed by blotting off the sample using a filter paper. The grid was then touched on a first drop of freshly prepared 0.75% (w/v) uranyl formate stain and immediately blotted off, followed by staining for 30 s on a second drop of stain. Imaging of the negatively stained samples were performed on a FEI Tecnai G2 12 Twin TEM (LaB6) operating at 120 kV and equipped with a Gatan CCD camera (4k x 4k) at 30,000x magnification. Data processing of the collected micrographs for the individual samples were performed with Relion 3.1.2 (Zivanov et al., 2020). Approximately 10,000 particles were autopicked using the gaussian blob picker within Relion and the extracted particles were subjected to reference free 2D classification.

### Cryo-EM sample preparation and data acquisition

Quantifoil holey carbon grids (Cu or Au, R2/1 or R2/2) were glow discharged for 45 s with a Glocube (Quorum technologies Ltd, UK). 3 μl of the complex was dispensed on the glow discharged grid, blotted for 3 s with a Whatman paper filter no. 1 at 10 °C and maintained at 90% humidity and then plunge frozen into liquid ethane (−180 °C) using a Leica GP plunger (Leica Microsystems, Austria).

For C5aR1pp-βarr1-Fab30 complex, cryo-EM data collection was performed on R2/2 Cu 300 mesh grid using a Titan Krios electron microscope (Thermofisher Scientific, USA) operating at 300 kV equipped with the Gatan Energy Filter. Movies were recorded in counting mode with a Gatan K2 Summit DED (Gatan, USA) using the automated SerialEM software at a nominal magnification of 165,000x and a pixel size of 0.82 Å at sample level. 6212 movie stacks consisting of 40 frames were recorded over a defocus range of 0.5 to 2.5 μm with a total dose of 49 e^-^/A^2^ and total exposure time of 5 s.

For CXCR4pp-βarr1-Fab30 complex, cryo-EM data collection was performed on R2/2 Cu 300 mesh grid using a Glacios electron microscope (Thermofisher Scientific, USA) operating at 200 kV. Movies were recorded in counting mode with a Gatan K3 DED (Gatan, USA) using the automated SerialEM software at a nominal magnification of 46,000x and a pixel size of 0.878 Å at sample level. 5,637 movie stacks consisting of 40 frames were recorded over a defocus range of 0.5 to 2.5 μm with a total dose of 49.3 e^-^/A^2^ and total exposure time of 2.9 s.

For V2Rpp-βarr2-Fab30 complex, cryo-EM data collection was performed on R2/2 Au 200 mesh grid using a Titan Krios electron microscope (Thermofisher Scientific, USA) operating at 300kV equipped with the Gatan Energy Filter. Movies were recorded in counting mode with a Gatan K2 Summit DED (Gatan, USA) using the automated SerialEM software at a nominal magnification of 165,000x and a pixel size of 0.82 Å at sample level. 9,720 movie stacks consisting of 40 frames were recorded over a defocus range of 0.5 to 2.5 μm with a total dose of 48.7 e^-^/A^2^ and total exposure time of 4 s.

For C5aR1pp-βarr2-Fab30 complex, cryo-EM data collection was performed on R2/2 Cu 300 mesh grid using a Glacios electron microscope (Thermofisher Scientific, USA) operating at 200 kV. Movies were recorded in counting mode with a Gatan K3 DED (Gatan, USA) using the automated SerialEM software at a nominal magnification of 46,000x and a pixel size of 0.878 Å at sample level. 8,614 movie stacks consisting of 40 frames were recorded over a defocus range of 0.5 to 2.5 μm with a total dose of 51 e^-^/A^2^ and total exposure time of 3 s.

### Cryo-EM data processing and model building

All image processing steps were performed in cryoSPARC version 3.3.2(Punjani et al., 2017) unless otherwise stated. In brief, for the C5aR1pp-βarr1-Fab30 complex, 6,212 movie stacks were subjected to Patch motion correction (multi), followed by CTF refinement with Patch CTF multi. 5,790 motion corrected micrographs with CTF fit resolution better than 4.5 Å were selected for further processing. 4,304,237 particle projections were automatically picked with blob picker, extracted with a box size of 480 pixels and fourier cropped to 64 pixels. The particle stack so obtained was subjected to multiple rounds of 2D classification. The class averages with clear secondary structural features were selected and re-extracted with a box size of 480 pixels and fourier cropped to 256 pixels resulting in a pixel size of 1.5375 Å. 295,922 re-extracted particles were then subjected to Ab-initio reconstruction and 3D classification/Heterogeneous refinement with C1 symmetry yielding 4 models. 84,655 particles corresponding to a dimer and containing 47.9% of the total particles were subjected to non-uniform refinement with C2 symmetry to yield a map with an estimated resolution of 3.41 Å (voxel size of 1.5375Å). Local resolution of all reconstructions was estimated using the Blocres within cryoSPARC version 3.3.2.

For the CXCR4pp-barr1-Fab30 complex data set, 5,637 movies were motion corrected using a patch of 5×5 patch within Patch motion correction (multi). Following CTF estimation, 5,236 motion corrected micrographs with CTF fit resolution better than 6Å were curated for further processing. 3,236,193 particles were automatically picked using the blob-picker sub-program and subsequently extracted with a box size of 480 pixels and fourier cropped to 64 pixels. The extracted particles were subjected to several rounds of 2D classification to remove junk particles. 104,707 particles corresponding to the clean class averages were selected, re-extracted with a box size of 480 pixels and fourier cropped to 256 pixels (pixel size of 1.65) and used to produce two ab-initio models. The particles corresponding to the two ab-initio models were subjected to heterogenous refinement/3D classification which produced a 3D class with clear dimeric conformation and a particle count of 53,387. This particle set was re-extracted with full box size of 480 pixels (pixel size of 0.878) and subjected to non-uniform refinement with C2 symmetry which converge to a map with 4.81Å resolution as estimated using the gold standard Fourier Shell Correlation (GFSC) using the 0.143 criterion.

For the V2Rpp-βarr2-Fab30 complex, 9,720 movies were motion corrected with 5×5 patches followed by CTF estimation with Patch CTF (multi). Following CTF refinement, 8,295 movies with CTF fit resolution better than 4.5Å were used for further processing. Particle picking from the curated micrographs was performed automatically with the blob picker sub-program to obtain an initial stack of 2,444,407 particles. The particles were then extracted with a box size of 512 pixels and fourier cropped to a box size of 64 pixels. The extracted particles were subjected to several rounds of reference free 2D classification. 2D class averages with evident secondary features containing 32,397 particles were extracted with a box size of 512 pixels and fourier cropped to a box size of 256 (pixel size of 1.64). This sub-set of particles was used for ab-initio reconstruction and subsequent rounds of 3D/Heterogeneous classification with C1 symmetry to obtain 2 models. 18,492 particles corresponding to a trimer were re-extracted with full box size of 512 pixels which refined to an overall resolution of 4.18 Å (voxel size of 0.82 Å) with NU refinement (C3 symmetry) according to the gold standard Fourier shell correlation (FSC) criterion of 0.143.

For the C5aR1pp-βarr2-Fab30 complex data set, 8,614 movies were motion corrected using Patch motion correction (multi) and subsequent CTF estimation was performed through Patch CTF (multi). 8,157 micrographs with CTF fit resolution better than 6Å were curated for particle picking using the blob picker sub-program. 4,012,616 particles were automatically picked and extracted with a box size of 512 pixels and fourier cropped to 64 pixels. Reasonable class averages after several rounds of reference free 2D classification yielded a particle set containing 54,587 particle projections, which was re-extracted with a box size of 512 pixels and fourier cropped to 360 pixels (pixel size of 1.2487Å) for subsequent used for ab-initio reconstruction generating two ab-initio models. Following heterogenous refinement/3D classification, the 3D class with evident features of a trimer and containing 38,206 particles was subjected to non-uniform refinement with C3 symmetry to yield a reconstruction at 4.41Å (final voxel size of 1.2487Å) as determined by gold standard Fourier Shell Correlation (FSC) using the 0.143 criterion.

### Model building and refinement

Coordinates from a previously solved IP6 bound βarr2 structure (PDB 5TV1) was used to dock the model into the EM density map of D6Rpp-βarr2-Fab30 using Chimera(Pettersen et al., 2004). The EM map was then used for manual rebuilding of the βarr1 residues and placing the phosphopeptide in COOT(Emsley et al., 2010). The rebuilt model was subjected to real space refinement in Phenix(Liebschner et al., 2019) to obtain a model with 94.77% of the residues in most favored region and 5.23% in the allowed region of the Ramachandran plot. The crystal structure of active βarr1 bound to vasopressin V2 receptor phosphopeptide (PDB 4JQI) was used as an initial model for model building and refinement against the C5aR1pp bound βarr1 density map. The model was docked into the coulombic map using Chimera, followed by manual rebuilding in COOT, and refinement of the rebuilt model with real space refinement in Phenix. The final refined model had 97.23% residues in the most favored region, while 2.77% in the allowed region of the Ramachandran plot.

The protomeric structure from the D6Rpp-βarr2-Fab30 (PDB 8GO9) complex solved in this study was used as an initial model to dock into the density map of V2Rpp-βarr2-Fab30 complex and regenerate the trimeric complex with C3 symmetry. The rigid body fitted trimeric model and the phosphopeptides were then rebuilt manually into the EM density map. The rebuilt trimeric coordinates with the phosphopeptides were subsequently subjected to real space refinement in Phenix to reach a final model with 95.05% in the favored region and 4.76% in the allowed region of the Ramachandran plot.

For model building into the 4.4Å C5aR1pp-βarr2-Fab30 coulombic map, the co-ordinates corresponding to V2Rpp peptide were deleted from the trimeric co-ordinates of V2Rpp-βarr2-Fab30 complex (PDB 8GOC), and the resulting model was docked into the EM map in Chimera. The “all atom refine” sub-module within the “refine” module in COOT was used for initial fitting of the model into the EM map, followed by manual rebuilding of the phosphopeptides. Multiple rounds of Phenix real space refinement combined with iterative model building yielded a model with 94.9% of the residues residing in the most favored region of the Ramachandran plot.

The dimeric co-ordinates from the cryo-EM structure of C5aR1pp-barr1-Fab30 (PDB 8GO8) without the phosphopeptide was used as an initial model to dock into the CXCR4pp-barr1-Fab30 EM map using Chimera. The docked model along with the coulombic map were imported into COOT and the model was subjected to “all atom refine” for fitting the atoms into the density. The phosphopeptide was manually built into the density to yield a complete model, which was subsequently used to refine the model against the EM map with Phenix real space refinement. The final refined model had 96.62% residues in the most favored regions and 3.38% in the allowed regions of the Ramachandran plot.

All the refined models were validated using “Comprehensive Validation (cryo-EM)” sub-module in Phenix. 3D reconstruction and model refinement statistics are provided as Supplementary Table S1. Figures in the manuscript have been prepared with Chimera(Pettersen et al., 2004) and ChimeraX(Pettersen et al., 2021) software. Domain rotation analysis was performed with PyMOL (Schrödinger, 2020).

### NanoBiT assay for βarr2^WT^ and βarr2^DM^ recruitment

βarr2^WT^ and βarr2^DM^ recruitment downstream of V2R and C5aR1 in response to AVP and C5a, respectively, was measured using NanoBiT (Enzyme linked complementation-based assay) assay following the protocol described earlier(Kawakami et al., 2022). Receptor constructs were tagged with SmBiT at the carboxyl-terminus, and βarr2 constructs were N-terminally tagged with LgBiT. Briefly, HEK293 cells were transfected with indicated receptor constructs (3.5 μg) and βarr2 (βarr2^WT/DM^) constructs (3.5 μg) using polyethylenimine (PEI) linear (Polysciences, Cat no. 19850) at a ratio of 1: 3 (DNA: PEI linear). After 16-18 h of transfection, cells were trypsinized, harvested, and resuspended in assay buffer (1XHBSS, 5 mM HEPES, pH 7.4, 0.01% BSA) containing 10 μM coelenterazine (GoldBio, Cat no. CZ2.5). Resuspended cells were seeded in a white flat bottom 96-well plate (100 μl well^-1^). After 2 h of incubation (90 min at 37 °C and 30 min at room temperature), basal luminescence was recorded using a multimode plate reader (FLUOstar Omega, BMG Labtech). Later, cells were stimulated with varying doses of indicated ligands followed by measurement of luminescence signal for 20 cycles. For data analysis, average of 5^th^ to 10^th^ cycle readings were taken and normalized with the lowest ligand dose signal, and fold normalized data was plotted using GraphPad Prism 9 software.

### NanoBiT assay for βarr trafficking

Agonist-induced βarr2^WT^ and βarr2^DM^ endosomal trafficking downstream of the receptors mentioned above was studied using NanoBiT assay as described in the recruitment experiment. The only exception from the recruitment assay was that the receptor constructs were not tagged with SmBiT, but rather βarr2 (βarr2^WT/DM^) and FYVE constructs N-terminally fused with SmBiT and LgBiT respectively were used for enzyme complementation. For each experiment, 3 μg of indicated receptors, 2 μg of SmBiT-βarr2^WT/DM^, and 5 μg of LgBiT-FYVE were used.

### NanoBiT assay for Ib30 reactivity

To assess Ib30 reactivity in response to an agonist for the mentioned receptors, nanobit assay was used following the same protocol as discussed in the βarr2^WT^ and βarr2^DM^ recruitment assay(Dwivedi-Agnihotri et al., 2022). For enzyme complementation, N-terminally SmBiT fused βarr1 and N-terminally LgBiT fused Ib30 were used. For transfection, 3 μg receptor except for CXCR7 (5 μg of which is transfected), 2 μg SmBiT-βarr1, and 5 μg LgBiT-Ib30 were used. Transfected cells were stimulated with varying doses of respective ligands (mentioned in corresponding figures).

### Receptor surface expression

Receptor surface expression in various assays was measured using a previously described whole cell-based surface ELISA assay^34^. To study the surface expression of the receptor, cells transfected with a particular receptor were seeded into a 0.01% poly-D-Lysine pre-coated 24-well plate at a density of 2×10^5^ cells well^-1^. Post 24 h of seeding cells were washed once with ice-cold 1XTBS, fixed with 4% PFA (w/v in 1XTBS) on ice for 20 min, washed again three times with 1XTBS, and blocked with 1% BSA (prepared in 1XTBS) at room temperature for 1.5 h. Afterward, cells were incubated with anti-FLAG M2-HRP antibody at 1: 5000 dilution (Sigma, Cat no. A8592) for 1.5 h, which was followed by three washes in 1% BSA. Subsequently, incubated with TMB-ELISA substrate (Thermo Fisher Scientific, Cat no. 34028) until a light blue color appeared. To quench the reaction, 100 μl of the colored solution was transferred to another 96-well plate containing 100 μl of 1 M H_2_SO_4_, and the absorbance was measured at 450 nm. Afterward, the TMB substrate was removed, washed twice with 1XTBS, and incubated with 0.2% (w/v) Janus Green (Sigma; Cat no. 201677) for 15 min at room temperature. Later, cells were washed with water to remove the excess stain, followed by the addition of 800 μl of 0.5 N HCl in each well. Thereupon, the colored solution was transferred to a 96-well plate for measuring the absorbance at 595 nm. The signal intensity was normalized by calculating the ratio of A450/A595 values followed by quantifying fold increase with respect to the A450/A595 value of negative control (mock transfection) and plotted using the GraphPad Prism (v9).

