## Supplemental Material for "Structural snapshots uncover a lock-and-key type conserved activation mechanism of β-arrestins by GPCRs"

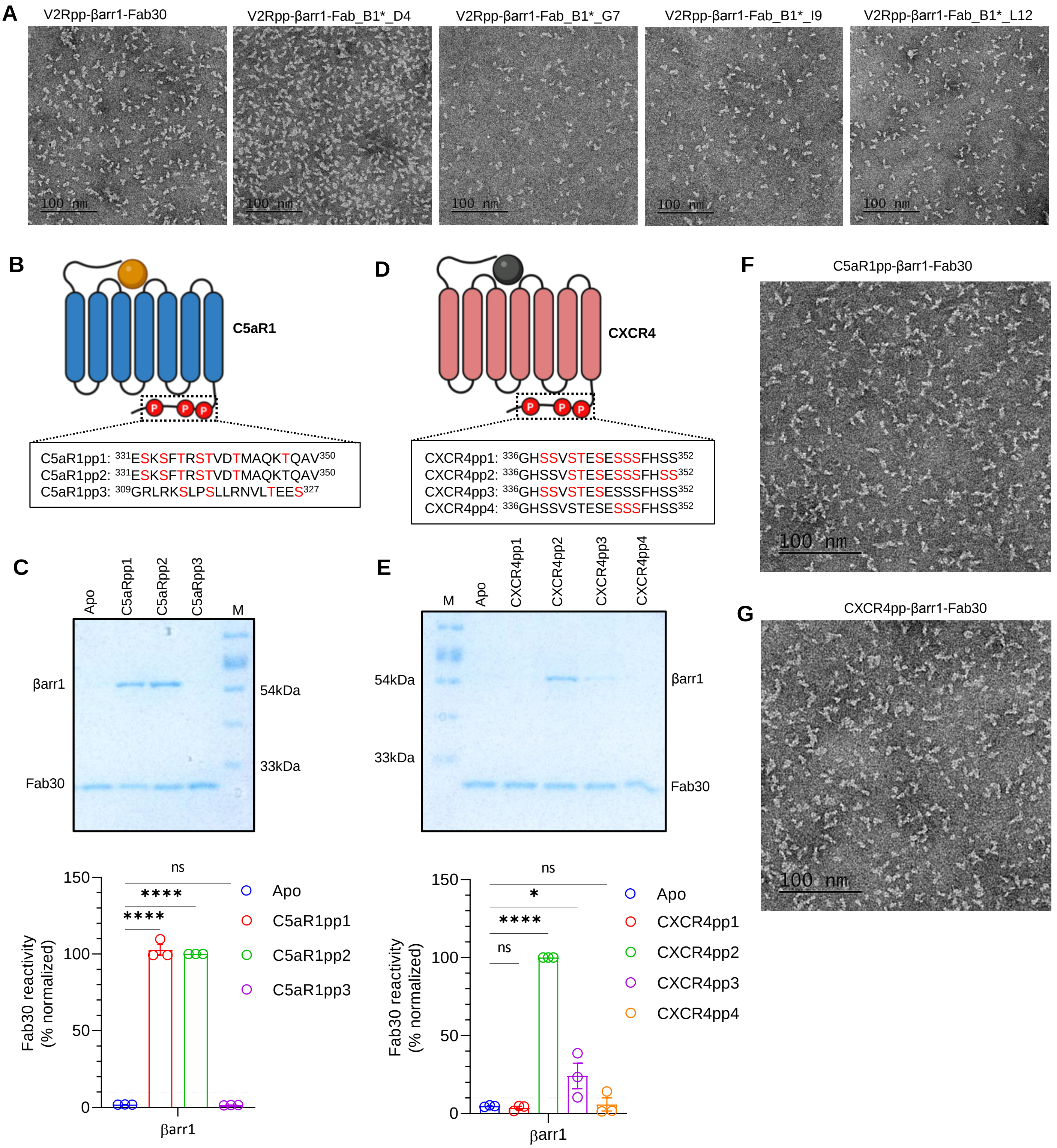

A
V2Rpp-βarr1-Fab30
V2Rpp-βarr1-Fab_B1*_D4
V2Rpp-βarr1-Fab_B1*_G7
V2Rpp-βarr1-Fab_B1*_I9
V2Rpp-βarr1-Fab_B1*_L12
100 nm
B
C5aR1
C5aR1pp1: 331ESKSFTRSTVDTMAQKTQAV350
C5aR1pp2: 331ESKSFTRSTVDTMAQKTQAV350
C5aR1pp3: 309GRLRKSLPSLLRNVLTEES327
D
CXCR4
CXCR4pp1: 336GHSSVSTESESSSFHSS352
CXCR4pp2: 336GHSSVSTESESSSFHSS352
CXCR4pp3: 336GHSSVSTESESSSFHSS352
CXCR4pp4: 336GHSSVSTESESSSFHSS352
F
C5aR1pp-βarr1-Fab30
G
CXCR4pp-βarr1-Fab30
C
Apo
C5aRpp1
C5aRpp2
C5aRpp3
M
βarr1
Fab30
54kDa
33kDa
E
M
Apo
CXCR4pp1
CXCR4pp2
CXCR4pp3
CXCR4pp4
54kDa
33kDa
βarr1
Fab30
Fab30 reactivity (% normalized)
ns
****
Apo
C5aR1pp1
C5aR1pp2
C5aR1pp3
βarr1
ns
*
****
Apo
CXCR4pp1
CXCR4pp2
CXCR4pp3
CXCR4pp4

**Figure S1. Reconstitution and characterization of V2Rpp/C5aR1pp/CXCR4pp-βarr1-Fab30 complexes.**

**(A)** Representative negatively stained micrographs of V2Rpp activated βarr1 with Fab30, Fab_B1*_D4, Fab_B1*_G7, Fab_B1*_I9 and Fab_B1*_L12. **(B)** Designed phosphopeptides corresponding to C5aR1. **(C)** Comparison of C5aR1pps in terms of Fab30 reactivity to βarr1 studied by co-immunoprecipitation. Densitometry-based quantification (mean ± SEM; n=3; normalized with respect to C5aR1pp2 signal as 100% (Ordinary one-way ANOVA, Tukey's multiple comparisons test). The exact *p* values are as follows: Apo vs. C5aR1pp1 ($p < 0.0001$), Apo vs. C5aR1pp2 ($p < 0.0001$), Apo vs. C5aR1pp3 ($p = 0.9982$). Source data are provided as a Source Data file (****$p < 0.0001$, ns = non-significant). **(D)** Designed phosphopeptides corresponding to CXCR4. **(E)** Comparison of CXCR4pps in terms of Fab30 reactivity to βarr1 studied by co-immunoprecipitation. Densitometry-based quantification (mean ± SEM; n=3; normalized with respect to CXCR4pp2 signal as 100% (Ordinary one-way ANOVA, Tukey's multiple comparisons test). The exact *p* values are as follows: Apo vs. CXCR4pp1 ($p = 0.9996$), Apo vs. CXCR4pp2 ($p < 0.0001$), Apo vs. CXCR4pp3 ($p = 0.0488$), Apo vs. CXCR4pp4 ($p = 0.9997$). Source data are provided as a Source Data file (*$p < 0.05$, ****$p < 0.0001$, ns = non-significant) (**F & G),** Representative negatively stained micrographs of C5aR1pp and CXCR4pp activated βarr1 with Fab30 respectively.

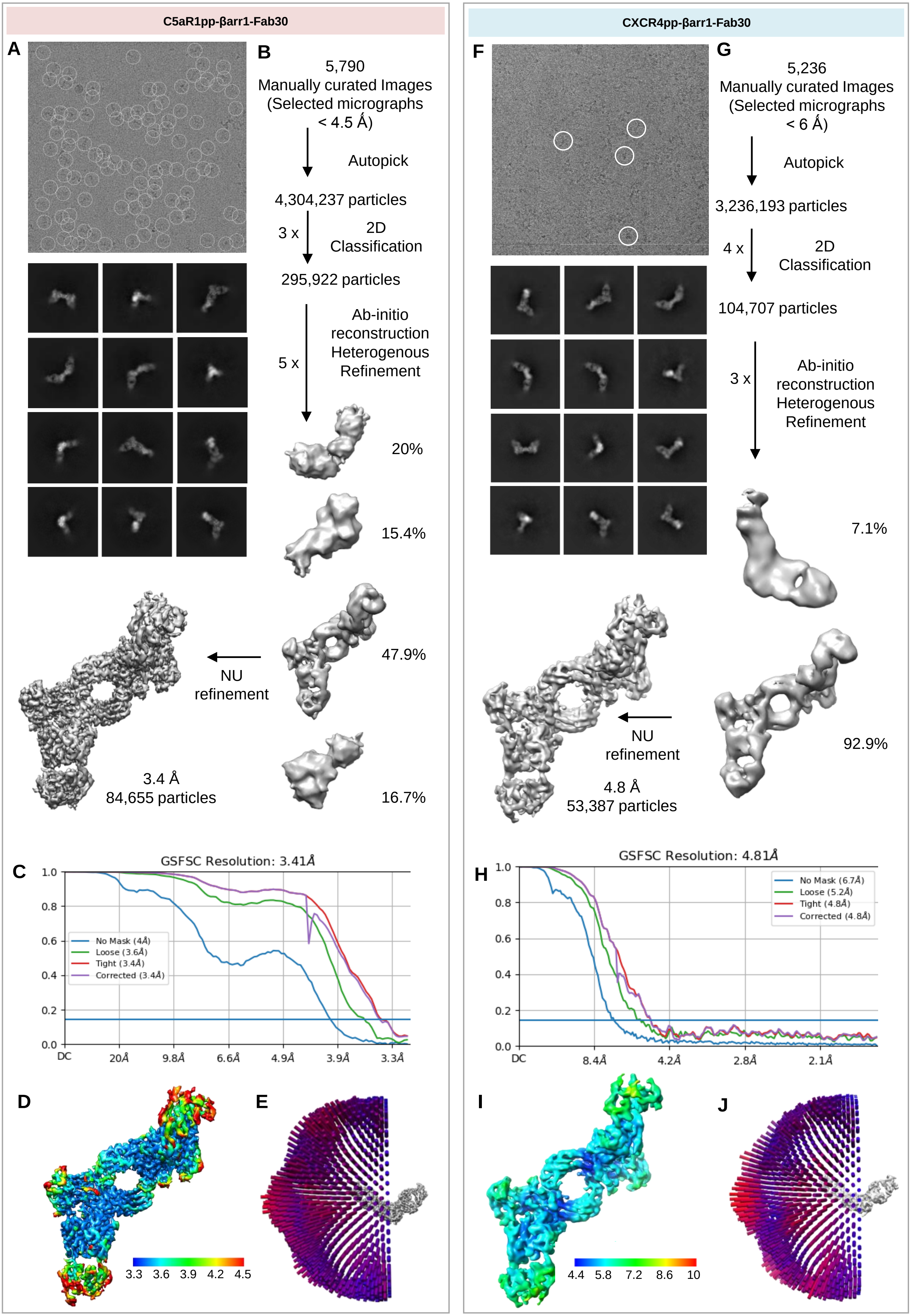

C5aR1pp-βarr1-Fab30
A
B
5,790
Manually curated Images
(Selected micrographs
< 4.5 Å)
Autopick
4,304,237 particles
3 x
2D
Classification
295,922 particles
5 x
Ab-initio
reconstruction
Heterogenous
Refinement
20%
15.4%
47.9%
16.7%
NU
refinement
3.4 Å
84,655 particles
C
GSFSC Resolution: 3.41Å
No Mask (4Å)
Loose (3.6Å)
Tight (3.4Å)
Corrected (3.4Å)
DC
20Å
9.8Å
6.6Å
4.9Å
3.9Å
3.3Å
D
E
3.3
3.6
3.9
4.2
4.5
CXCR4pp-βarr1-Fab30
F
G
5,236
Manually curated Images
(Selected micrographs
< 6 Å)
Autopick
3,236,193 particles
4 x
2D
Classification
104,707 particles
3 x
Ab-initio
reconstruction
Heterogenous
Refinement
7.1%
92.9%
NU
refinement
4.8 Å
53,387 particles
H
GSFSC Resolution: 4.81Å
No Mask (6.7Å)
Loose (5.2Å)
Tight (4.8Å)
Corrected (4.8Å)
DC
8.4Å
4.2Å
2.8Å
2.1Å
I
J
4.4
5.8
7.2
8.6
10

### Figure S2. Cryo-EM reconstruction of C5aR1pp-βarr1-Fab30 complex and CXCR4pp-βarr1-Fab30.

**(A)** Representative motion corrected micrograph (top) and selected 2D class averages of cryo-EM particle images of C5aR1pp-βarr1-Fab30 complex (bottom). **(B)** Schematic cryo-EM data processing workflow. **(C)** Gold standard fourier shell correlation curve (GFSC). Gold standard threshold of 0.143 was used to determine the overall resolution of the map. **(D)** Local resolution map of the 3D reconstruction in front view. **(E)** Angular plot of the particles used for 3D reconstruction of C5aR1pp-βarr1-Fab30. **(F)** Representative motion corrected micrograph (top) and selected 2D class averages of cryo-EM particle projections of CXCR4pp-βarr1-Fab30 complex (bottom). **(G)** Schematic cryo-EM data processing workflow. **(H)** Gold standard fourier shell correlation curve (GFSC) at 0.143 threshold. **(I)** Local resolution map of the 3D reconstruction in front view. **(J)** Angular plot of the particles used for 3D reconstruction of CXCR4pp-βarr1-Fab30.

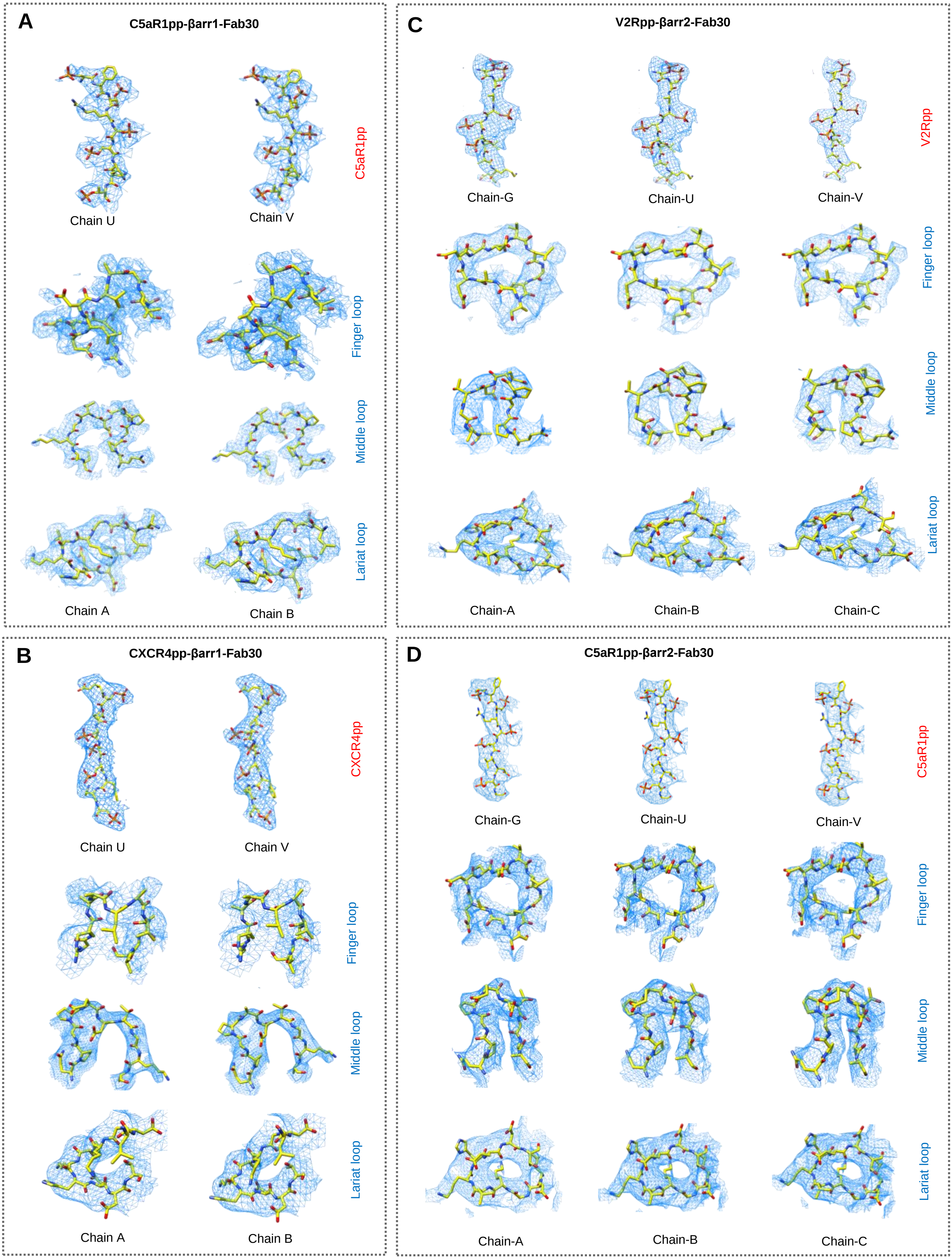

A
C5aR1pp-βarr1-Fab30
C5aR1pp
Chain U
Chain V
Finger loop
Middle loop
Lariat loop
Chain A
Chain B
C
V2Rpp-βarr2-Fab30
V2Rpp
Chain-G
Chain-U
Chain-V
Finger loop
Middle loop
Lariat loop
Chain-A
Chain-B
Chain-C
B
CXCR4pp-βarr1-Fab30
CXCR4pp
Chain U
Chain V
Finger loop
Middle loop
Lariat loop
Chain A
Chain B
D
C5aR1pp-βarr2-Fab30
C5aR1pp
Chain-G
Chain-U
Chain-V
Finger loop
Middle loop
Lariat loop
Chain-A
Chain-B
Chain-C

### Figure S3. Exemplary electron density maps.

EM densities for the phosphopeptides, finger loops, middle loops and lariat loops in the **(A)** C5aR1pp activated βarr1 **(B)** CXCR4pp activated βarr1 **(C)** V2Rpp activated βarr2 **(D)** C5aR1pp activated βarr2.

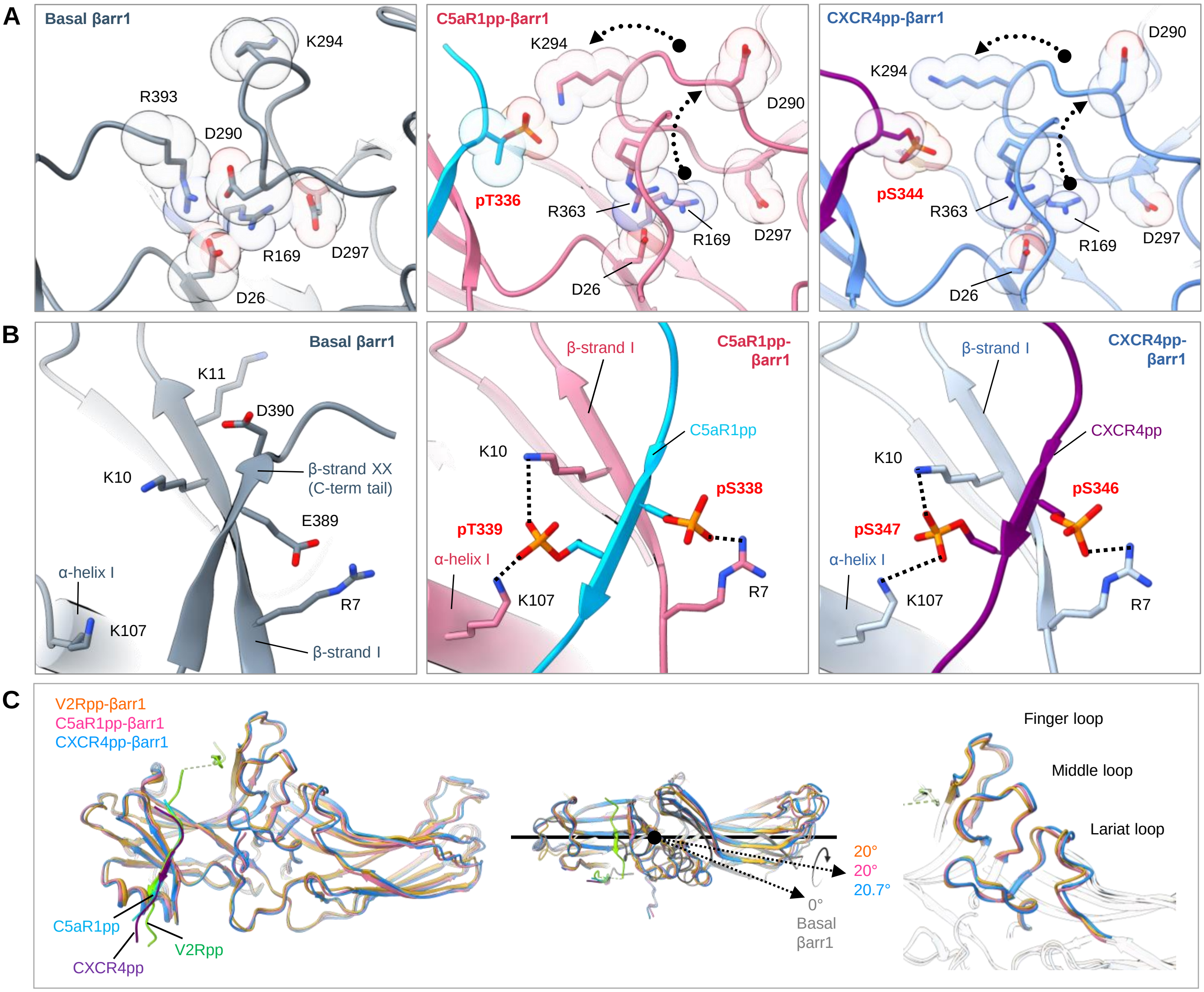


**Figure S4. Conformational features of activated βarr1 bound to C5aR1pp and CXCR4pp.**

**(A)** Polar core environment in basal βarr1 (PDB 1G4M, left) and disruption of polar core interactions upon binding of C5aR1pp (middle) and CXCR4pp to βarr1 (right). A new interacting residue $R^{363}$ of βarr1 C-terminal tail engages with $D^{26}$ of the polar core. **(B)** Three element interaction network consisting of βarr1 C-terminal β-strand XX, α-helix1 and β-strand1 in the basal state of βarr1 (left). Binding of the phosphopeptides C5aR1pp or CXCR4pp to βarr1 results in the displacement of the β-strand XX, and engages the phosphopeptide C5aR1pp (middle) and CXCR4pp (right) into the N-domain groove of βarr1 through hydrogen bonds and polar interactions. **(C)** Overall structural features of V2Rpp-βarr1-Fab30 compared to C5aR1pp and CXCR4pp bound βarr1 (left). Domain rotation of βarr1 bound to V2Rpp, C5aR1pp and CXCR4pp show similar C domain rotation (~20°C) (middle). Superposition of key loops in the three structures show minimal changes in conformations (right).

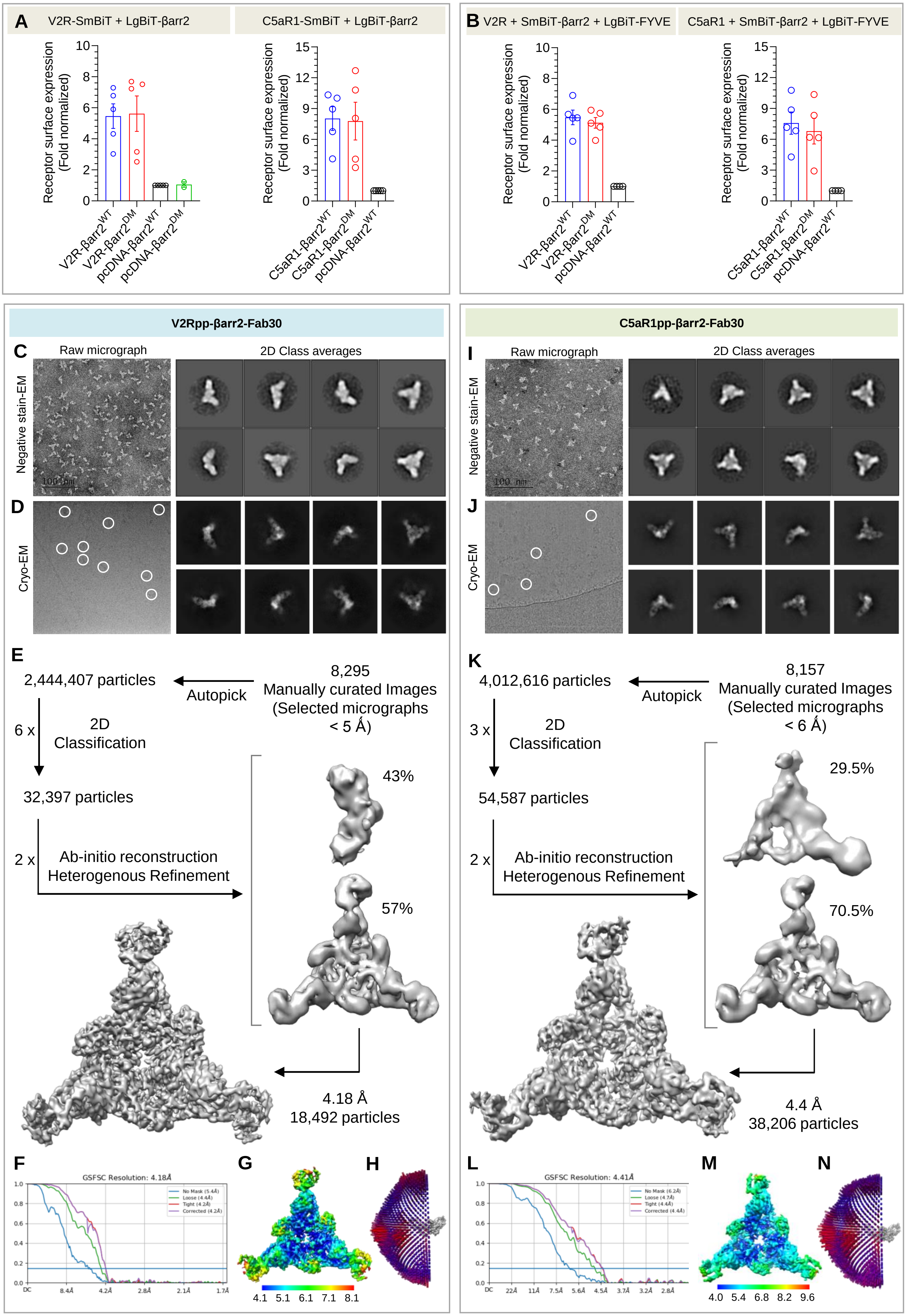

A
V2R-SmBiT + LgBiT-βarr2
C5aR1-SmBiT + LgBiT-βarr2
Receptor surface expression (Fold normalized)
V2R-βarr2WT
V2R-βarr2DM
pcDNA-βarr2WT
pcDNA-βarr2DM
C5aR1-βarr2WT
C5aR1-βarr2DM
B
V2R + SmBiT-βarr2 + LgBiT-FYVE
C5aR1 + SmBiT-βarr2 + LgBiT-FYVE
V2Rpp-βarr2-Fab30
C5aR1pp-βarr2-Fab30
C
Raw micrograph
2D Class averages
Negative stain-EM
100 nm
D
Cryo-EM
I
J
E
2,444,407 particles
Autopick
8,295
Manually curated Images
(Selected micrographs
< 5 Å)
6 x
2D
Classification
32,397 particles
2 x
Ab-initio reconstruction
Heterogenous Refinement
43%
57%
4.18 Å
18,492 particles
K
4,012,616 particles
8,157
Manually curated Images
(Selected micrographs
< 6 Å)
3 x
54,587 particles
29.5%
70.5%
4.4 Å
38,206 particles
F
GSFSC Resolution: 4.18Å
No Mask (5.4Å)
Loose (4.4Å)
Tight (4.2Å)
Corrected (4.2Å)
G
4.1 5.1 6.1 7.1 8.1
H
L
GSFSC Resolution: 4.41Å
No Mask (6.2Å)
Loose (4.7Å)
Tight (4.4Å)
Corrected (4.4Å)
M
4.0 5.4 6.8 8.2 9.6
N

**Figure S5. Surface expression and cryo-EM data processing workflow of V2Rpp-βarr2-Fab30 and C5aR1pp-βarr2-Fab30.**

**(A & B)** Surface expression of indicated receptors in the $\beta arr2^{WT}$ and $\beta arr2^{DM}$ recruitment and endosomal trafficking assays were measured by using whole cell ELISA (mean±SEM; n=4). **(C)** Representative micrograph and 2D class averages of negatively stained sample of V2Rpp-βarr2-Fab30 complex. **(D)** Representative cryo-EM motion corrected micrograph and selected 2D class averages. **(E)** Data processing workflow for 3D reconstruction of the V2Rpp-βarr2-Fab30 complex. **(F)** Fourier shell correlation curve (GFSC) of the final 3D reconstruction at standard threshold of 0.143 indicates a global resolution of 4.18Å. **(G)** Local resolution map of the 3D reconstruction. **(H)** Angular plot of particles used for final reconstruction of V2Rpp-βarr2-Fab30 complex. **(I)** Representative micrograph and 2D class averages of negatively stained sample of C5aR1pp-βarr2-Fab30 complex. **(J)** Representative cryo-EM motion corrected micrograph and selected 2D class averages. **(K)** Data processing workflow for 3D reconstruction of the C5aR1pp-βarr2-Fab30 complex. **(L)** Fourier shell correlation curve (GFSC) of the final 3D reconstruction at standard threshold of 0.143 indicates a global resolution of 4.41Å. **(M)** Local resolution map of the 3D reconstruction. **(N)** Angular plot of particles used for final reconstruction of C5aR1pp-βarr2-Fab30 complex.

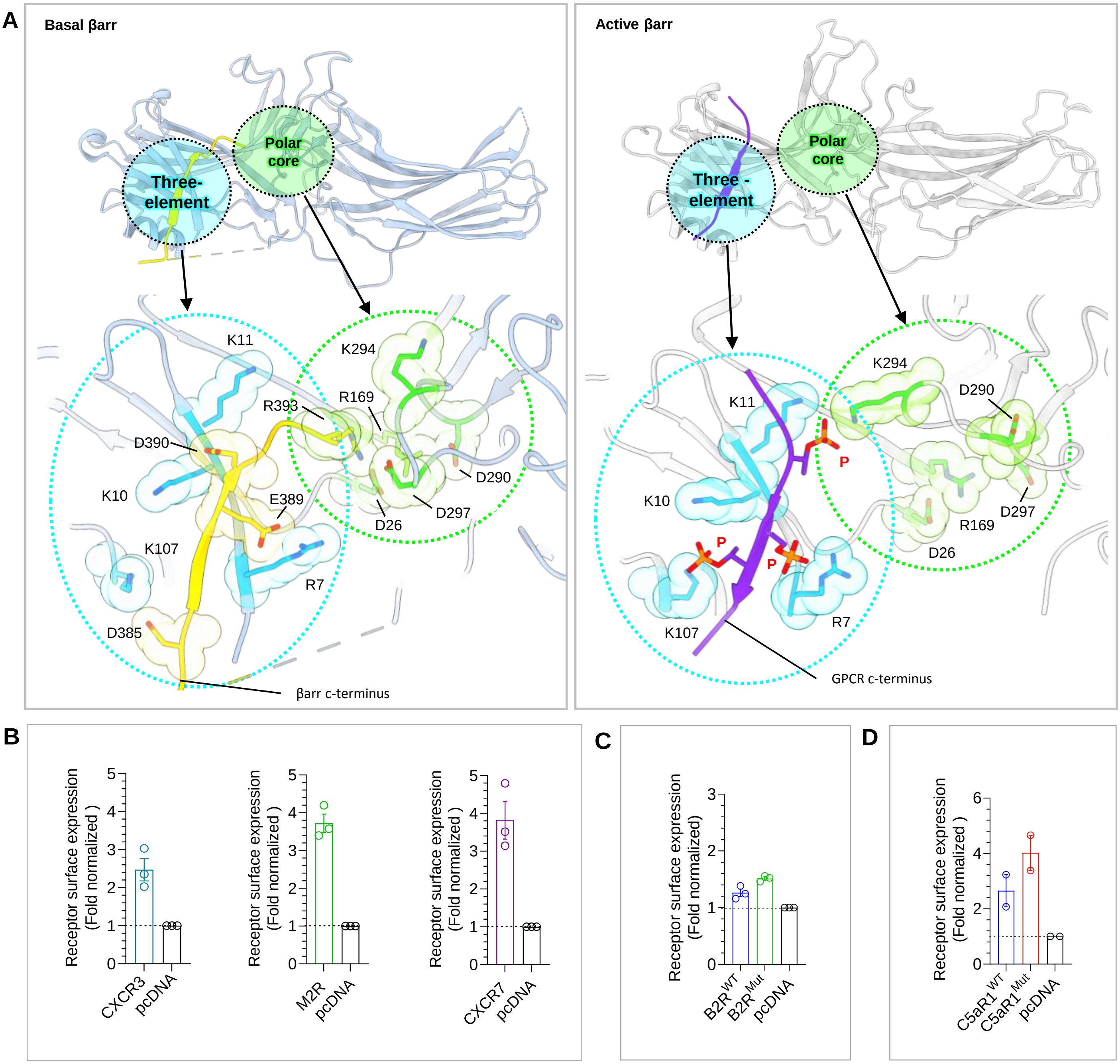


**Figure S6. Structural activation of βarrs by P-X-P-P and surface expression profiles of GPCRs for functional validation experiments.**

(**A**) Phosphoresidues in P-X-P-P pattern directly contact with the structural elements; "the polar core network" and "the three element interaction" in β-arrestins thereby maintaining and stabilizing the activated state. **(B, C & D)** Surface expression of indicated receptors in the Ib30 reactivity assay was measured by using whole cell ELISA (mean±SEM; n=3 for CXCR3, M2R, CXCR7, and B2R constructs; n=2 for C5aR1$^{WT}$ and C5aR1$^{Mut}$

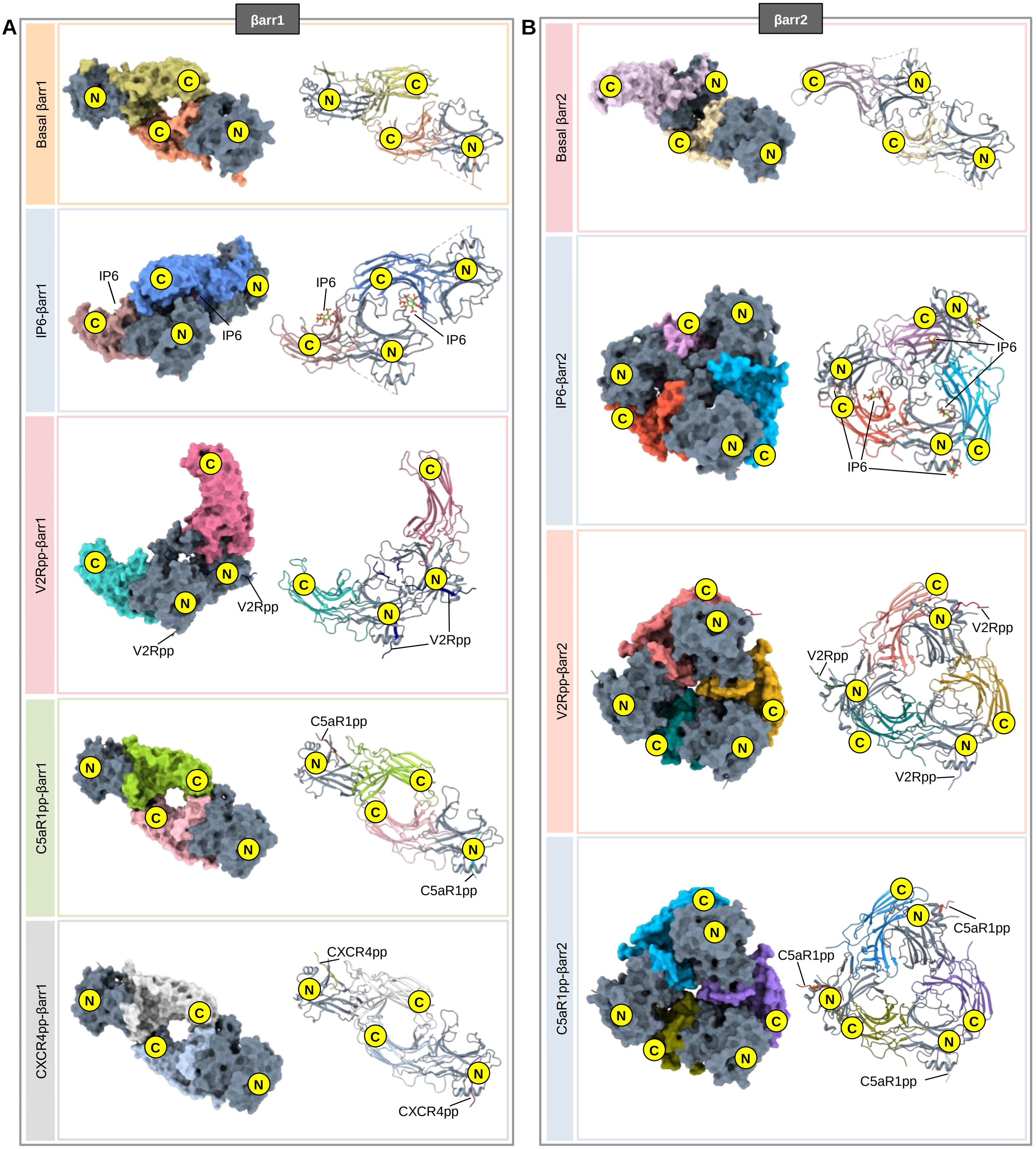


**Figure S7. Oligomerization states of βarr1 and βarr2.**

**(A)** Oligomerization mode, intermolecular interfaces and domain organization of different βarr1 oligomers. The biological assemblies for the previously solved crystal structures of basal βarr1 (PDB 1G4M), IP6-βarr1 (PDB 1ZSH) and V2Rpp bound βarr1 (PDB 4JQI) along with the cryo-EM structures solved in this study have been shown for comparison. **(B)** Comparative analysis of the intermolecular interfaces and oligomerization mode of βarr2 oligomers. The biological assemblies for the previously solved crystal structures of basal βarr2 (PDB 3P2D) and IP6-βarr2 (PDB 5TV1) along with the cryo-EM structures solved in this study have been shown here for comparison. N-domains have been colored in gray, while C-domains have been represented in various colors.
